# Resolvin D-series regulates Phospholipase D both during inflammation and resolution by modulating phagocyte functions

**DOI:** 10.1101/827295

**Authors:** Ramya Ganesan, Karen M. Henkels, Krushangi Shah, Xavier de la Rosa, Stephania Libreros, Nagarjuna R Cheemarla, Charles N. Serhan, Julian Gomez-Cambronero

## Abstract

A successful acute inflammatory response results in the elimination of infectious agents by neutrophils and monocytes, followed by resolution and repair by tissue-resident and recruited macrophages. D-series resolvins are pro-resolving mediators involved in resolution and tissue repair, their intracellular signaling mechanism(s) are of interest. Here, we report that D-series resolvins activate phospholipase D (PLD), a ubiquitously expressed membrane lipase in modulating phagocyte functions. The mechanism for PLD-mediated actions of RvD5 in polarizing macrophages (M1-M2) was found to be two-pronged: *(a)* RvD5 inhibits post-transcriptional modifications, by miRs and 3’exonulceases that process PLD2 mRNA, thus increasing PLD2 expression and activity; *(b)* RvD5 enhances PLD2-S6K signaling and Actin expression required for membrane expansion and efferocytosis. In addition, investigating RvD5’s actions in second organ reflow injury, we found that RvD5 did not reduce lung neutrophil myeloperoxidase levels in PLD2^-/-^ mice compared to WT and PLD1^-/-^ mice, pointing to a unique role of PLD2 as the relevant isoform in RvD5-mediated resolution. These results demonstrate that RvD5-PLD2 are attractive targets for therapeutic interventions in vascular inflammation such as I/R injury and cardiovascular diseases.

## INTRODUCTION

Inflammation is a response initiated by harmful stimuli, such as infections or injury that stimulate endogenous mediators (1). These mediators immediately generate an inflammatory exudate locally, which then enters circulation and activates first line of defense mechanism, infiltrating plasma proteins and innate inflammatory cells such as neutrophils, monocytes and macrophages by extravasation to the site of inflammatory insult (such as infection or injury) (2). Macrophages are also key players in inflammatory diseases and are the phagocytic cells involved, among other processes, in the clearance of apoptotic neutrophils after an injury known as the “resolution phase” (3).

Macrophages play a critical role in maintaining tissue homeostasis (4-6) in pathologies such as ischemia/reperfusion injury, tumor microenvironment, obesity, and conditions of chronic inflammation (7-11). Their diverse functions, including cytokine production and phagocytosis, place macrophages at the balance of pro-inflammation and resolution, both necessary components of the healing process (5,6). Macrophages can polarize into M1 or M2 phenotypes (4,5). M1 cells are regarded as generally pro-inflammatory and have a high microbicidal capacity and secrete pro-inflammatory cytokines, such as interleukin-1β (IL-1b), interleukin-12 (IL-12) and tumor necrosis factor α (TNF-α) (7). In contrast, M2 cells mediate resolution of inflammation by secreting interleukin-10 (IL-10) and transforming growth factor β (TGF-β) (7). Pro-inflammatory signals, such as toll-like receptor (TLR), ligands and interferon γ (IFNγ), induce polarization to the M1 phenotype, while anti-inflammatory signals, such as interleukin-4 (IL-4) and IL-10, induce polarization to the M2 phenotype (9). Macrophage class switch is critical to proper macrophage function and tissue homeostasis. Disruption of the M1/M2 balance results in various pathologies (5,7,12).

The process of inflammation has two phases, wherein after an inflammatory insult the body initiates a pro-inflammatory response, which eliminates infectious agents, followed resolution and repair, an important role of tissue-resident macrophages. Resolution, the process of returning to baseline conditions occurs via a well-regulated anti-inflammatory program aimed to restore homeostasis (13). Resolvins and protectins, are families of lipid mediators that have crucial roles in the resolution of inflammation, including the initiation of tissue repair (13,14). Resolvins below to the new family of specialized lipid mediators derived from omega-3 essential fatty acids, docosahexanoic acid (DHA) and eicosapentanoic acid (EPA) that are known to have both pro-resolution and anti-inflammatory actions (15,16). For example, Resolvin E1 exhibits a time-dependent response to inflammation resolution. During the short or acute inflammatory response period (0-4h), resolvins promote phagocytosis via activation of S6Kinase that phosphorylates ribosomal S6 protein resulting in phagosome formation and hence phagocytosis (17). Resolvins and protectins both stimulate innate killing mechanisms to manage bacterial loads and stimulate clearance of bacteria (3). Resolvin D5 activates a specific receptor denoted human GPR32 (18). However, the intracellular mechanism of downstream signaling in macrophages is of interest.

Phospholipase D (PLD) is a ubiquitously expressed membrane-associated lipase that hydrolyzes phosphatidylcholine (PC) into free choline and phosphatidic acid (PA) (19-21). PLD is upregulated in response to various cell stressors, such as hypoxia and nutrient starvation (22,23). Moreover, PLD2^-/-^ mice produced less pro-inflammatory cytokines in a sepsis model (24). The product of the PLD reaction, PA, is itself a mitogen and a critical secondary messenger that activates many downstream pathways leading to cell growth and proliferation, vesicle trafficking and cell migration (19-21). Additionally, PA is conical in shape and carries a negative charge, meaning its accumulation in the membrane results in membrane curvature needed for cell migration (23,25-27). Although PLD/PA play a role in macrophage adhesion (25,28), no role of PLD in macrophage polarization has been described as of yet.

Phospholipase D (PLD, with the two major mammalian isoforms, PLD1 and PLD2) has a pivotal role in cellular activities such as chemotaxis and phagocytosis (21,29-34). Aberrant PLD expression and activity is implicated in inflammation (23,35-38) and other cellular functions (21,39). PLD is also actively involved in pro-inflammatory cytokine recruitment, reactive oxygen species (ROS) generation, chemotaxis, and cell invasion (40-43). In this report, we set out to understand the role of resolvins in inflammation and resolution involving PLD as an intracellular signaling molecule.

## RESULTS

### PLD expression and activity in macrophages

Since well-appreciated mechanisms separating self-limited acute inflammation from delayed resolution exists (3,13), we questioned whether PLD would play a role in the more delayed response, specifically in the resolution of inflammation. In order to study the role of PLD in inflammation and resolution, it was critical to measure the expression levels of PLD in the different macrophage subpopulations that occur in the inflammation-resolution process. As such, we polarized differentiated peripheral blood M0 human macrophages into either M1 (pro-inflammatory) or M2 (anti-inflammatory) phenotype by cytokine treatment and conducted a series of experiments to establish the basal levels and authenticity of those populations (*Supplemental* ***Fig. S1A-E***). **Fig. S1A** shows the characteristic morphologies of M0, M1 and M2 macrophages; ***Fig. S1B*** presents the gene expression of markers of differentiation, with iNOS2 being predominately expressed in M1 and Arg1 and PPARγ predominately expressed in M2 macrophages. We have previously shown that PLD regulates the PPAR family of transcription factors (44). Endogenous protein expression levels of iNOS2 and Arg1 in M1 and M2 macrophages are presented in Western blots (***Fig. S1C***), confirming gene expression data. We also tested the functionality of macrophages. M1 had quantitatively higher capacity in phagocytosis of *E. coli* than M2 macrophages (***Fig. S1D***); conversely M2 had quantitatively higher capacity in efferocytosis of apoptotic neutrophils than M1 macrophages (***Fig. S1E***), results that were in agreement with (45,46).

Concentrating on PLD, we measured basal gene expression for *Pld1* and *Pld2* from M0, M1 and M2 macrophages (***Fig. S1F***); basal levels of protein (***Fig. S1G***) and endogenous lipase activity from M0, M1 and M2 macrophages (***Fig. S1H***). M1 macrophages had a level of basal activity higher than the M0 controls. PLD1 gene and protein were highly expressed in M2 macrophages, whereas PLD2 gene and protein were highly expressed in both M1 and M2 macrophages. Our interest was to investigate why both PLD1 and PLD2 are present in M2 macrophages as such high levels.

### D series Resolvins impact PLD’s gene expression and enzyme activity in macrophages

Having observed the basal expression of PLD1 and PLD2 in M1 and M2 macrophages, we next determined if Resolvins, the specialized pro-resolving mediators (SPMs), had any effect on PLD mRNA levels in macrophages. We focused our interest on Resolvins of the D series (RvDs), including RvD1, RvD2, RvD3, RvD4 and RvD5, and their effect on PLD activity of differentiated macrophages. We treated M0, M1 and M2 macrophages with vehicle (0.1% ethanol) or 10 nM D-series Resolvins (RvD1-5).

S*upplemental* ***Figs. S2***, ***S3*** show the effects of different periods of incubation of Resolvins-D1-5 (at 10 nM) on *Pld1* (***Fig. S2***) and *Pld2* (***Fig. S3***) gene expression in M0, M1 or M2 macrophages. We observed that even though RvD1 or RvD3 had some punctual effect on M2’s on specific conditions, both RvD4 and RvD5 are the Resolvins that more consistently had the largest effect in altering PLD gene expression. Since 6 hr of incubations of Resolvins with macrophages was not enough to elicit significant changes in PLD gene expression, we concentrate in **Fig. 1** on 24 hr of incubation with macrophages. There are statistically significant increases of *Pld1* in M1 (**Fig. 1A**) and M2 macrophages (**Fig. 1**A). Interestingly, RvD5 decreased *Pld2* gene expression in M1 macrophages (**Fig. 1C**), whereas RvD5 induced a robust increase in Pld2 in M2 (**Fig. 1D**). In fact, in M2 macrophages other resolvins (RvD1, RvD3 and RvD4) were also able to induce an increase in *Pld2* gene expression, albeit at lower quantitative levels than RvD5.

**Figure 1.**
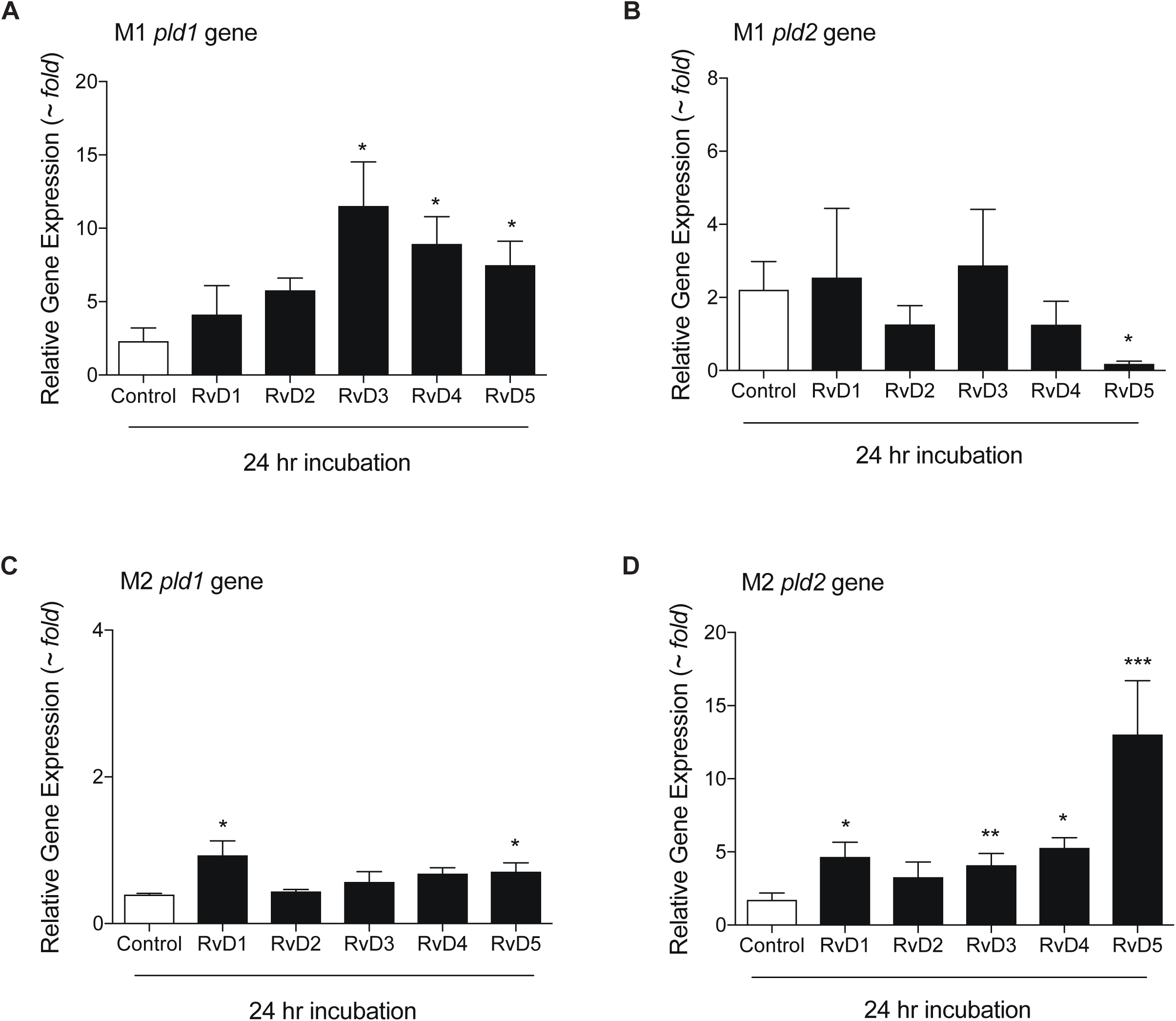
D-series Resolvins actions on PLD1 and PLD2 gene expression in M1 and M2 macrophages. Effect of D-series Resolvins (each at 10 nM) on endogenous PLD1 (A,B) or PLD2 (C,D) gene expression in M1 or M2 macrophages. For each condition, 1×10^6^ cells were treated with 10 nM of each of the indicated Resolvins-D series for 24 hours in continuous culture. Gene expression was measured by qRT-PCR. Data presented are means ± SEM (n>3 independent experiments); statistical significance (*p < 0.05, **p < 0.01, and ***p < 0.001) was evaluated with one-way ANOVA and Tukey’s post hoc comparing samples with controls (vehicle only) for each panel. See supplemental figures S2 and S3 for a full range of Pld1 and Pld2 gene changes at different time lengths of macrophages incubation with RvDs.

We then measured PLD enzyme activity at 6, 12, 18 or 24 hr for PLD activity in M0, M1 and M2 macrophages (**Fig. 2**). These time points were chosen since resolvins are involved in acute inflammation resolution and start early during inflammation soon after macrophages arrive to the site of inflammation. We observed the positive effects of RvD4 and RvD5 on PLD activity from M0, M1 and M2 macrophages was time-dependent (statistically significantly increased after 24 h incubation with RvD5). As observed earlier for PLD gene expression, RvD4 and RvD5 maximally stimulate PLD activity. It should also be noted that the *in vitro* PLD assay monitors both PLD1 and PLD2 combined activity. Since activity is upregulated by RvD4/5 and so is PLD1 gene expression (and PLD2 is downregulated) it seems plausible that the effects of RvD4/5 on M1 are mediated by PLD1. Conversely, since the main PLD isoform expressed in M2 is PLD2, it seems plausible that the effects of RvD4/5 on M2 are mediated by PLD2.

**Figure 2.**
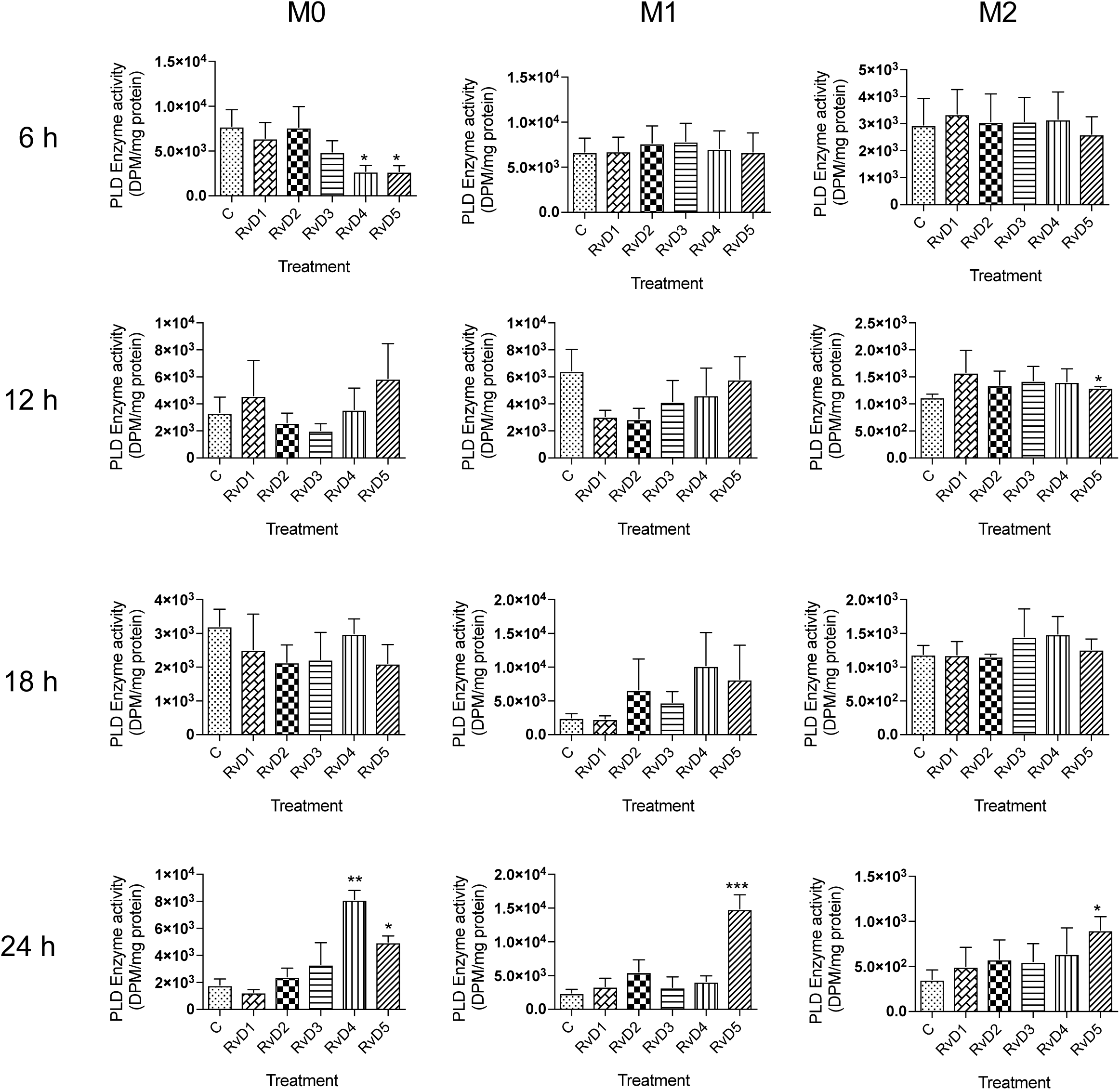
Actions of D-series Resolvins on PLD enzymatic activity in M0, M1 and M2 macrophages. Effect of D-series Resolvins (each at 10 nM) on total PLD activity in M0, M1, or M2 macrophages. For each condition, 1×10^6^ cells were treated with 10 nM of each of the indicated Resolvins-D series at 6,12,18 or 24 hours in continuous culture. Total PLD activity was measured by an enzyme assay using [3H]-butanol in an *in vitro* transphosphatidylation reaction to yield [3H]-phosphobutanol. Data presented are means ± SEM (n>3 independent experiments); statistical significance (*p < 0.05, **p < 0.01, and ***p < 0.001) was evaluated with one-way ANOVA and Tukey’s post hoc comparing samples with controls (vehicle only) for each panel.

We were interested in understanding this latter phenomenon and investigating other *Pld* genes. There are PLD enzyme isoforms other than PLD1 or PLD2, namely PLD3, PLD 4 and PLD 6. The scheme (*Supplemental* **Fig. *S4A***) shows the *“classical”* (PLD1 and PLD2) and *“non-classical”* (PLD3, PLD4 and PLD6), (note: PLD5 has not been fully described), where mammalian PLD architecture depends on the protein architecture regulatory domains. All PLD’s have one or two HKD “lipase signature” domains, but only PLD1 and PLD2 have significant domains (PX and PH) that allow the protein to anchor strongly to cellular or the intracellular membranes. In macrophages, we observed that not all levels of *Pld* genes are equally expressed. *Supplemental* **Fig. *S4B*** compares the basal gene expression levels of *Pld1, Pld2, Pld3, Pld4*, and *Pld6* in M1 and M2 macrophages, indicating that the levels of basal expression of *Pld1* and *Pld2* are the highest. The gene expression levels of *Plds* in M0, M1, and M2 macrophages treated with 10 nM series-D Resolvins at several lengths of time are shown in *supplemental* ***Fig. S5*** (for *Pld3*), ***Fig. S6*** (for *Pld4*), and ***Fig. S7*** (for *Pld6*), that indicates that RvD from 18 to 24 hr incubation changed the patterns of expression of all these three genes. **Fig. 3 (A-C)** shows that RvD5 causes inhibition of *Pld* gene expression in M1 macrophages and activation in M2 macrophages, but only *pld6* gene expression was inhibited by RvD2 in M1 macrophages. Overall, *Pld3, Pld4* and *Pld6* follow the same pattern of expression in response to RvD’s as *Pld2* (**Figs. 1B,1D**), which is, in M1 macrophages RvD5 inhibits and in M2 activates PLD gene expression. PLD1 has a different pattern from all the others genes in that RvD5 always activates M1 or M2 (**Figs. 1A,1C**). Thus, RvD5 contributes the most to the increased expression of PLD in anti-inflammatory/pro-resolving M2 macrophages, whereas PLD expression is reduced in inflammatory-M1 macrophages. This may mean that during resolution of inflammation, PLD levels are high via RvD5 regulation suggesting PLD’s novel role in inflammation-resolution. Because the levels of *Pld1* and *Pld2* are the highest expressed of all PLD’s studied (**Fig. 3B**) and they both used PC as substrate, we concentrated on PLD1 and PLD2 for the rest of this study.

**Figure 3.**
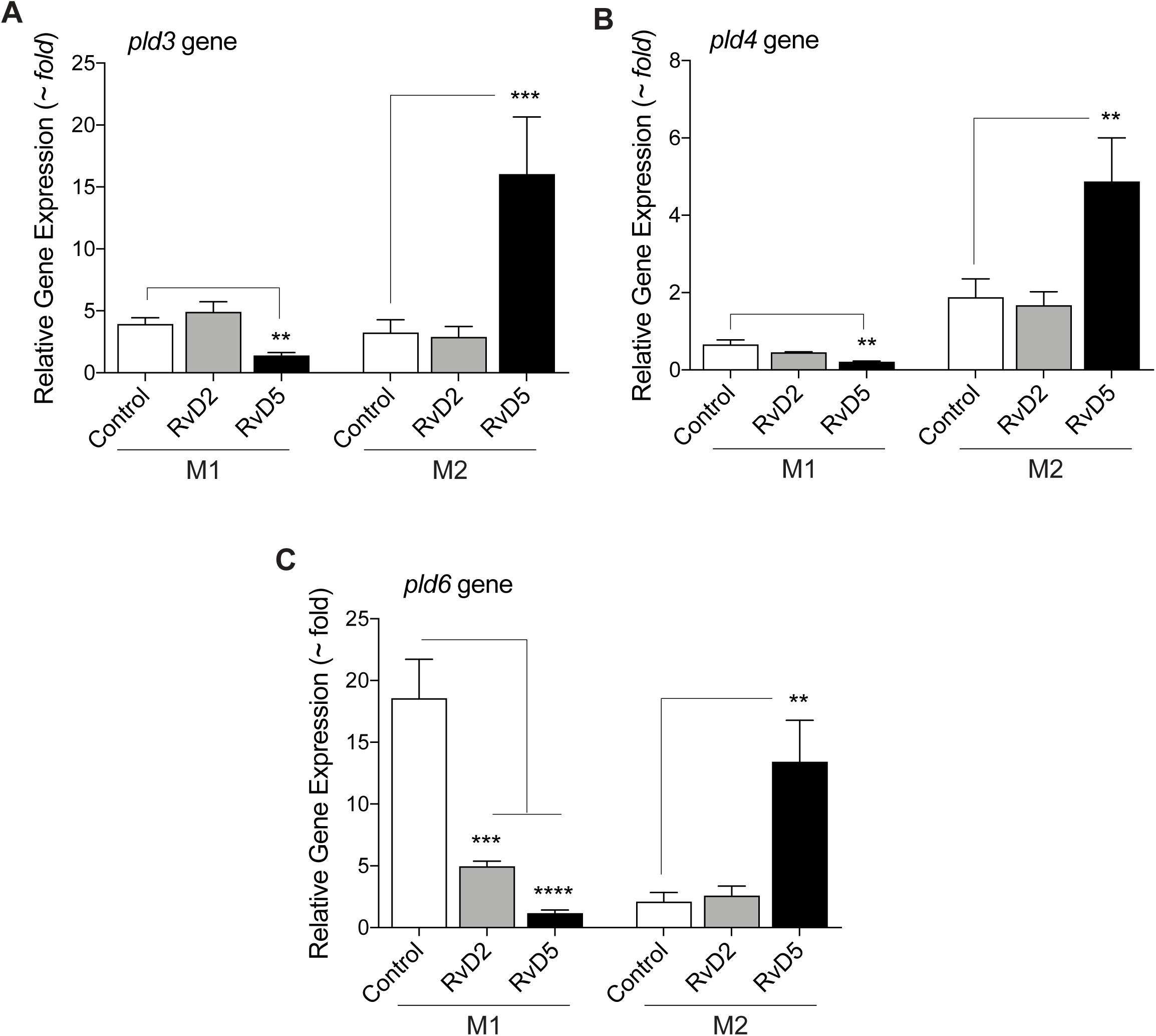
Actions of specific D-series Resolvins on “non-classical” PLD’s gene expression in M1 or M2 macrophages. (A-C) Gene expression levels for Pld3 (A), Pld4 (B), and Pld6 (C), measured by qRT-PCR in M1 or M2 macrophages treated with 10 nM RvD2 or RvD5 for 24 hours. Data presented are means ± SEM (n>3 independent experiments); statistical significance (*p < 0.05, **p < 0.01, and ***p < 0.001) was evaluated with one-way ANOVA and Tukey’spost hoc comparing conditions to the controls (vehicle) for each panel and each phenotype. See Supplementary Figures S5, S6 and S7 for a full range of Pld3, Pld4 and Pld6 gene changes at different time lengths of macrophage incubation with RvD’s.

### Resolvin D5 increases macrophage PLD protein expression and function

The actions of both RvD4 and RvD5 on PLD protein expression in M1 and M2 macrophages (at different lengths of time of incubation) are shown in *supplemental* ***Fig. S8A*** and corresponding densitometry quantification of protein bands in *supplemental* ***Figs. S8B-C***. PLD1 protein is stimulated with RvD4 but more so, quantitatively, by RvD5. PLD1 protein expression is stimulated at 6-24 hr whereas PLD2 is stimulated earlier (30 min-6 hr). These results, suggested that there could be post-transcriptional regulation of PLD levels by resolvins.

To investigate the functional relevance of this consistent action of RvD5 on PLD, we performed macrophage functional assays using human macrophages that were silenced for PLD1 or PLD2 and subjected to RvD5 treatment during the duration of the assays. We found that silencing PLD1 or PLD2 statistically significantly inhibited RvD5’s action on phagocytosis of *E. coli* by M1 macrophages (**Fig. 4A**) whereas, silencing PLD2 only affected RvD5’s action on phagocytosis of apoptotic neutrophils (efferocytosis) by M2 macrophages (**Fig. 4B**). Thus, PLD is important for RvD5-mediated functions on macrophage phagocytosis, with the PLD2 isoform being more involved functionally than the PLD1 isoform.

**Figure 4.**
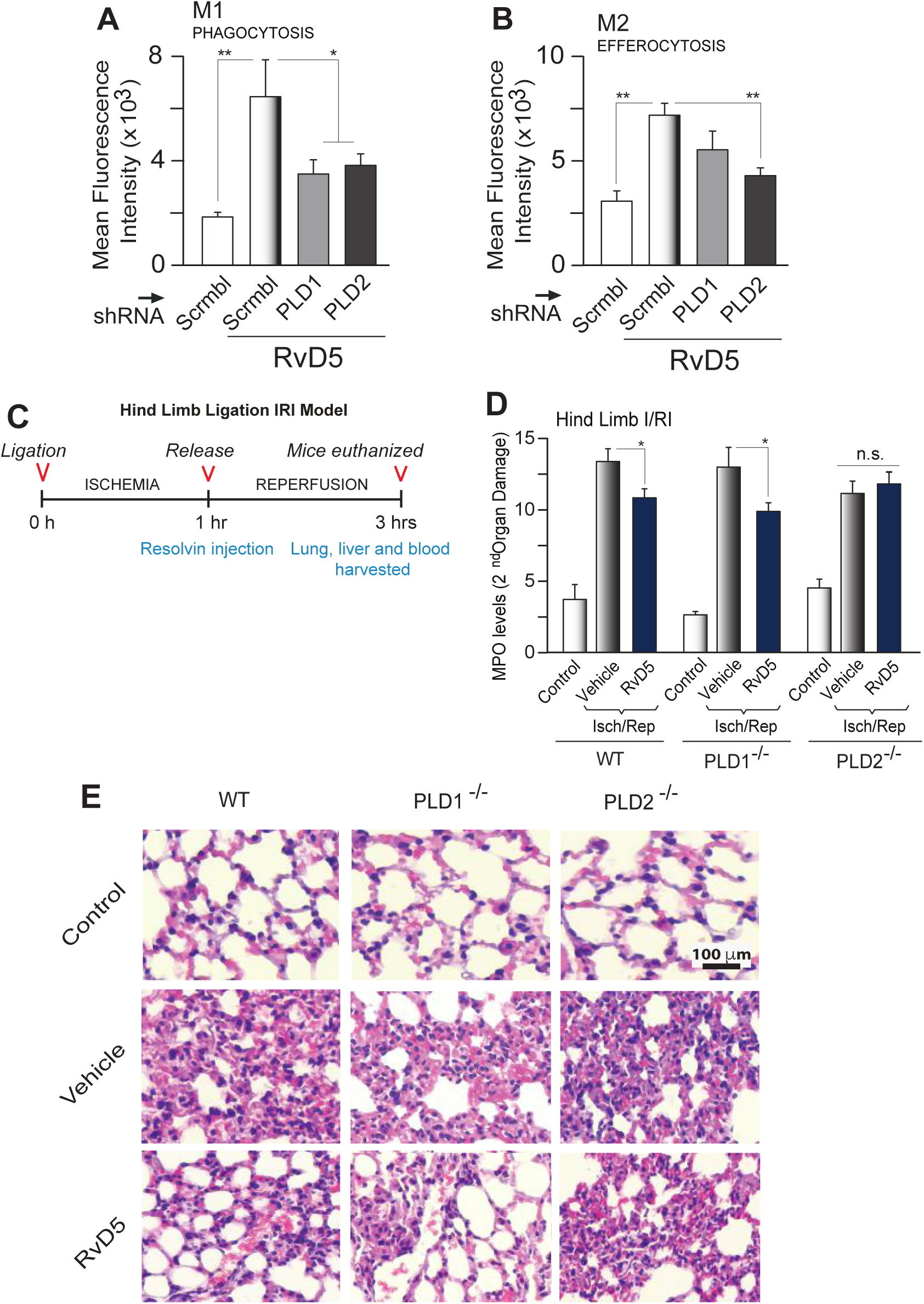
RvD5 protects mice from second organ injury in a PLD dependent manner. (A) Phagocytosis. Fluorescence intensity of phagocytosis of BacLightGFP-labeled E. coli by real-time microscopy of M1 macrophages upon silencing PLD1 or PLD2 with RNAi for 72 hr, followed by incubation with 10 nM RvD5 for 1 hr. *E. coli* was added in a ratio of 1:50 to 3×105 macrophages. (B) Efferocytosis. Fluorescence intensity of fluorescent-(CellTraceTM-CFDA)-labeled apoptotic neutrophils (PMNs) was measured by flow cytometry, using M2 macrophages upon silencing PLD1 or PLD2 with RNAi for 72 hr, followed by incubation for 1 hr with 10 nM RvD5. A macrophage to CFDA-labeled apoptotic PMN ratio was added at 1:5 with 3×105 cells/well. (C) Schematic timeline of the used hind limb ischemia/reperfusion injury (HLIR) procedure. (D) Myeloperoxidase (MPO) levels (units/μg (wet weight) of protein in tissue) were measured using ELISA from lung lysates of WT, PLD1-/- or PLD2-/-mice treated with vehicle or 10 nM of RvD5. (E) Representative photomicrographs (from a total of 10 fields/per condition observed) showing pathology of lung sections by H&E staining from control, vehicle or RvD5 treated mice. Scale bar=100 μm. Results are expressed as mean ± SEM from n>3 independent experiments. Statistical significance (*p < 0.05, **p < 0.01, and ***p < 0.001; n.s.=not significant) was evaluated with one-way ANOVA and Tukey’s post hoc comparing samples with controls (Scrbl=scrambled RNA in (A,B) or control in WT mice in (D)).

### RvD5 activates PLD-mediated inflammation resolution *in vivo*

To investigate the contribution and validate the *in vivo* relevance of the effects of RvD5-PLD signaling in sterile injury and inflammation, we performed ischemia reperfusion injury and assessed second-organ injury to the lungs (46). Our lab has demonstrated PLD’s role in cell migration in many cell types, including neutrophils and macrophages (27,35-37), in pathological conditions such as cancer and atherosclerosis (27,34). Uncontrolled inflammation is linked to defective generation of pro-resolving mediators, such as in asthma, which suggests that resolution of acute inflammation is critical (47). As the damaged, ischemic tissue initiates an intense innate immune response (inflammation), we hypothesized that administration of RvD5 to a second organ injury model of inflammation/injury mediated by PLD could keep this process at bay and yield a more favorable outcome. To test this hypothesis, we administered vehicle or RvD5 to PLD1^-/-^, PLD2^-/-^ and WT mice in a mouse model of second organ injury to the lungs using a model of sudden reperfusion after hind limb ischemia.

The outline for the hind limb ischemia/reperfusion injury (HLIR) procedure is in **Fig. 4C**. Hind limb ischemia was performed for 1 h at which time vehicle or 500 ng of RvD5 were administered by i.v. injection and then mice were reperfused for 2 h. Following completion of reperfusion, lungs were harvested and used for measuring myeloperoxidase (MPO) levels in the damaged lung tissues. Myeloperoxidase (MPO) levels were assessed using ELISA (**Fig. 4D**) of lung lysates from WT, PLD1^-/-^ or PLD2^-/-^ treated with vehicle or 500 ng RvD5 at the beginning of reperfusion. We observed that RvD5 reduced MPO levels in WT and PLD1^-/-^ mice, suggesting a reduction in PMNs infiltration, while RvD5 did not affect MPO levels in the PLD2^-/-^ mice. Based on these results, we conclude that RvD5 functions via the PLD2 isoform and not the PLD1 isoform.

We also performed H&E staining on the lung tissue sections to observe lung pathology (second-organ injury) after hind-limb ischemia reperfusion injury (**Fig. 4E**). We observed increased neutrophil infiltration and loss of honeycomb lattice structure in the WT, PLD1^-/-^ and PLD2^-/-^ mice after ischemia-reperfusion injury compared to non-ischemia controls. This lung injury and neutrophil accumulation were reduced in RvD5 treated mice in WT and PLD1^-/-^, but not in PLD2^-/-^ suggesting that RvD5 functions via PLD2 during HLIR. Based on these results, we conclude that PLD2 is important for RvD5-mediated effects on neutrophil infiltration *in vivo*.

### M1 to M2 macrophage polarization is induced by PLD

It is known that specific resolvins have a role in promoting a macrophage “class switch” toward M2-like phenotype (48,49) and the polarization that results in increased efferocytosis by M2 macrophages and promotes resolution of inflammation (45,46). Studying the role of PLD altering M1 and M2 specific marker expression in the naïve M0 macrophages as indicated in the previous figures, led us to further investigate the role of PLD in the process of inflammation-resolution. As known, after an inflammatory insult, macrophages polarize to M1 and later undergo transient polarization to M2 macrophages to promote anti-inflammation and resolution via efferocytosis (45,46). To study if PLD had a role in macrophage polarization during inflammation and later in resolution of inflammation, we undertook two separate types of experiments: *(a)* PLD silencing and transfection of naïve/non-polarized M0 macrophages; and *(b)* PLD silencing and transfection of M1 macrophages. **Fig. 5A,B** indicate that non-differentiated macrophages (M0) can be induced to express an M1-like phenotype with PLD ectopic expression; that is, overexpressed PLD2, statistically significantly increased M1 marker expression CD80 and decreased M2 marker expression CD206 (**Fig. 5A**) whereas silencing PLD2 gene increased M2 marker, CD206 expression (**Fig. 5B**). As controls for this and the next series of experiments, PLD activity upon either RNAi silencing or ectopic overexpression of PLD1 or PLD2 expression plasmids in M1 macrophages is shown in **Fig. 5C**. Thus, we found that PLD expression induces macrophage polarization, from M0-M1 and not M0-M2 confirming a role of PLD early in inflammation.

**Figure 5.**
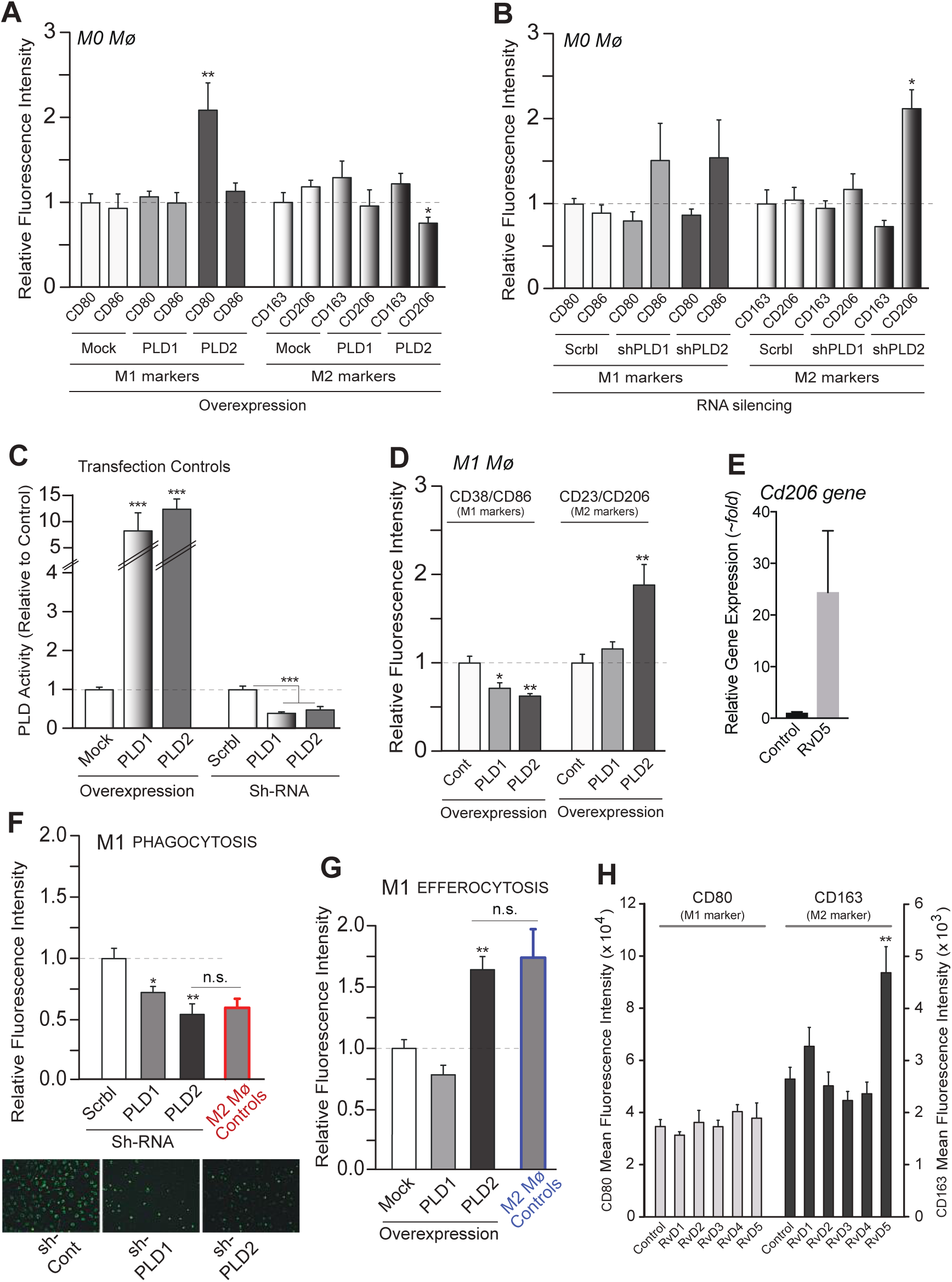
M1 to M2 macrophage reprogramming induced by PLD. (A,B) Flow cytometry analysis of surface expression of polarization markers for CD80 and CD86-positive cells (M1 markers) or CD163 and CD206-positive cells (M2 markers). Experiments were performed in M’zero’ (M0) macrophages overexpressing either PLD1 or PLD2 (1.5 ug DNA plasmid per condition) after 24 hr transfection (A) or silenced with shPLD1 or shPLD2 (100 nM per condition) after 72 hr transfection with shRNAs (“Scrbl”, scrambled RNA control) (B). Shown in (C) are the transfection controls to show efficiency of either overexpression or silencing of PLD1 and PLD2 plasmids or shRNA, respectively. (D) Flow cytometry analysis of surface expression of polarization markers for CD38/CD86-positive cells (M1 markers) or CD23/CD206-positive cells (M2 markers) in M1 macrophages upon ectopic overexpression of PLD1 or PLD2 plasmids. See Supplementary Figure S9 for flow cytometry plots. (E) Gene expression of CD206 using qRT-PCR with or without RvD5 (10 nM) treatment in culture for 24 hrs. (F) Phagocytosis of M1 macrophages silenced with PLD1 or PLD2 shRNA. Fluorescence intensity of phagocytosis of BacLightGFP-labeled E. coli was measured by real-time microscopy of M1 macrophages upon silencing PLD1 or PLD2 with 100 nM shRNA (“Scrbl”, scrambled RNA control) for 72 hr, followed by incubation with 10 nM RvD5 for 1 hr. A cell to E. coli ratio was 1:50 with 3×10^5^ cells. (inset=example of fluorescent-field view from the microscope). (G) Efferocytosis of M1 macrophages after transfection of PLD1 or PLD2 expression plasmids. Fluorescence intensity of green fluorescence intensity from efferocytosis of green fluorescence (CellTraceTM-CFDA)-labeled apoptotic neutrophils (PMNs) was measured by flow cytometry, upon transfection of PLD1 or PLD2 expression plasmids for 48 hr, followed by incubation for 1 hr with 10 nM RvD5. A macrophage to CFDA-labeled apoptotic PMN was added to a 1:5 ratio with 3×10‘ cells/well. (H) Effect of ResolvinsD1-5 on surface expression of polarization markers on M1 and M2 macrophages, measured by flow cytometry. Data in bar graphs are mean ± SEM (n>3 independent experiments; for real-time microscopy, sample size n∼60 cells read for each condition); statistical significance (*p < 0.05, **p < 0.01, and ***p < 0.001) was evaluated with one-way ANOVA and Tukey’s post hoc multiple comparisons.

Next, we began the experiments with M1 macrophages instead of M0. PLD is upregulated during inflammation, although to date, a phase and time specific role of PLD in late inflammation and resolution have not been studied. M1 macrophages were transfected with plasmids to overexpress PLD1 or PLD2 or with PLD silencing with RNAi. Flow cytometric analysis of CD38/CD86-positive cells (M1 markers) or CD23/CD206-positive cells (M2 markers) was then conducted (**Fig. 5D**). See gaiting strategy on *supplemental* ***Fig. S9***. Since CD206 is an M2 phenotype marker, we investigated *CD206* gene expression upon PLD2 overexpression in the presence of RvD5, wherein we found that RvD5 by itself induces the expression of CD206 (**Fig. 5E**). Additionally, transfected macrophages with PLD1 or PLD2 expression plasmids induced morphology changes in the cells that appear larger and more elongated (data not shown), which is a phenotype more consistent with the presence of M2-like cells.

We next tested whether PLD would alter macrophage functions. To investigate this, we silenced PLD1 or PLD2 and measured phagocytosis of *E. coli* by M1 macrophages (**Fig. 5F**). We observed that silencing PLD1 or PLD2 statistically significantly decreased macrophage phagocytic functions: there was a decrease in *E. coli* phagocytosis. The inset in **Fig. 5F** presents a visual control of cells silenced with either PLD1 or PLD2 siRNA. The red bar indicates the level of phagocytosis of M2 macrophages treated and measured in similar conditions as M1. When PLD2 was silenced in M1 macrophages, *E. coli* phagocytosis decreased. Thus, silencing PLD2 in macrophages lead to a decrease of phagocytosis, pointing at the importance of this molecule in leukocyte functionality.

We also measured efferocytosis, a known function of resolving macrophages, and found that overexpressing PLD1 and PLD2 statistically significantly augmented efferocytosis (**Fig. 5G**). The blue bar indicates the level of efferocytosis of M2 macrophages treated and measured in similar conditions as M1. Thus, when M1 macrophages were PLD2-overexpressed (becoming “M2-like induced”), apoptotic PMN phagocytosis (efferocytosis) increased. We found that PLD had a role in class-switch from M1 to M2 with the expected functions of those cell subpopulations.

Lastly, incubation of macrophages with RvD5 resulted in an elevated CD163 M2 marker but no changes on CD80 M1 marker when directly compared to control (**Fig. 5H**). RvD1 was recognized as an inducer of macrophage polarization (48-50) but such similar role for RvD5 has not been previously described. In summary, after finding out differential expression of PLD1 and PLD2 in the different polarized macrophage subpopulations, we observed that overexpressing PLD2 decreased M1 markers and increased M2 marker expression in M1 macrophages, and showed an increase in apoptotic PMN efferocytosis by these macrophages. Both of these results confirmed our earlier results, indicating a role of PLD in macrophage polarization and thus function.

### Mechanism for PLD-mediated effect of RvD5 in macrophages: *(a)* Involvement of miRs and PARN

To understand the possible mechanism(s) behind this newly uncovered role of PLD-mediated RvD5 signaling, we first analyzed two post-transcriptional mechanisms that could target PLD transcripts and could be altered by RvD5: miRs and PARN mRNA deadenylase.

Protein expression of PLD2 (normalized to GAPDH) derived from Western blot analysis in M0, M1 and M2 resting macrophages (*supplemental* ***Fig. S8A*** and **Fig. 6A**) shows that PLD2 protein is highly expressed in both M1 and M2 macrophages with respect to M0. There could be several reasons for this. We have studied previously post-translational modification of PLD2 transcripts and reported how that is under regulation of certain miRs (22,26). For macrophages, we selected miR-138 and miR-887 that are predicted to bind to the 3’UTR of PLD2 (**Fig. 6B**). The demonstration that transfected miR-138 and miR-887 decrease PLD2 expression in macrophages is shown in **Fig. 6C**. The levels of endogenous miR expression in resting macrophages in M0 are higher than in M1 or M2 macrophages (**Fig. 6D**), which could partially explain the results of protein expression (*supplemental* ***Fig. S8A***). Furthermore, levels of miR expression in response to 10 nM RvD5 are further reduced (**Fig. 6E**). Our miRNA results are in agreement with previous findings for a role of Resolvins on miR biology (51-55). In the present study, RvD5 causes the inhibition of the inhibitory properties of miRs, therefore, indirectly RvD5 stimulates PLD2 expression.

**Figure 6.**
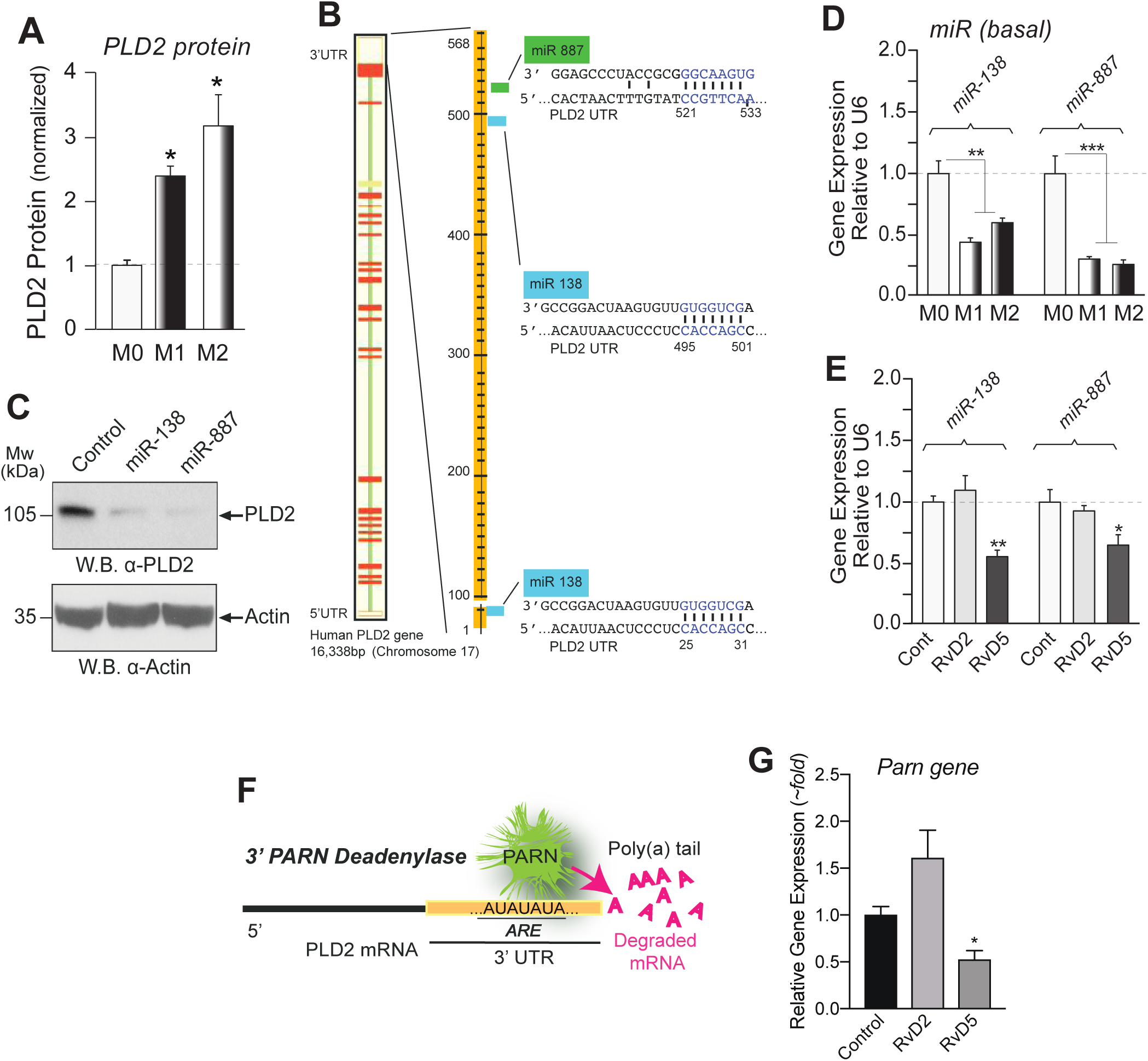
Mechanism for PLD-mediated actions of RvD5 in macrophages involving miRs and PARN. (A) Protein expression of PLD2 (normalized to GAPDH) derived from Western blot analysis in M0, M1 and M2 macrophages. Scheme depicting the 3’UTR of PLD2 with predicted target sites for miR-138 and miR-887. (C) Demonstration that transfected miR-138 and miR-887 decrease PLD2 expression in macrophages. (D) Levels of endogenous PLD-miR expression (by qRT-PCR) in M0, M1 or M2 macrophages. (E) Levels of PLD-miR expression from macrophages that had been incubated with or without 10 nM RvD2 or RvD5 for 6 hrs. (F) Scheme depicting the 3’ exonuclease activity of Poly(A) Ribonuclease (PARN) as cleaving the poly-A tail of PLD mRNA transcripts in the cell cytoplasm, leading to mRNA decay. (G) Effect of RvD2 or RvD5 on PARN gene expression. Data from RAW264.7 macrophages are represented as means ± SEM (n>3 independent experiments); statistical significance (*p < 0.05, **p < 0.01, and ***p < 0.001) was evaluated with one-way ANOVA and Tukey’s post hoc comparing samples with controls (vehicle only) for each panel.

There are other possible post-transcriptional ways to regulate PLD2 expression, for example; 3’exonulceases that can process mRNA transcripts in the cytoplasm, thus elevating the rate of mRNA decay (56). The scheme presented in **Fig. 6F** depicts the 3’ exonuclease activity of Poly(A) Ribonuclease (PARN) as cleaving the poly-A tail of mRNA transcripts leading to the ultimate destruction of mRNA by exosomes and P-bodies. Incubation of macrophages with RvD5 decreases the basal level of *Parn* gene expression (**Fig. 6G**). PARN silencing increases *pld2* gene expression (56), an effect that is also observed upon treatment with RvD5. Therefore, inhibition of an inhibitory PLD2 pathway (PARN) results in overall activation of PLD2, initiated by RvD5.

### Mechanism for PLD-mediated effect of RvD5 in macrophages: *(b)* Involvement of S6K and Actin

To further understand the mechanism(s) behind the newly uncovered role of PLD-mediated RvD5 signaling, we next analyzed two intracellular pathways in which PLD effectors are implicated, S6K and Actin polymerization, and asked if they could be altered by RvD5. S6K has been established as a morphogenic protein (57) through Actin activation and it is involved in cell shape change and is part of the underlying mechanism that regulates cytoskeletal structures needed for phagocytosis. Basal S6K gene expression is highest in M2 macrophages (**Fig. 7A**). We observed a statistically significant positive effect on S6K mRNA levels following RvD4 and especially in RvD5 treatment in M2 macrophages (**Fig. 7B**). This is similar to PLD2 upregulation by RvD5 in M2’s described earlier (**Fig. 1D**).

**Figure 7.**
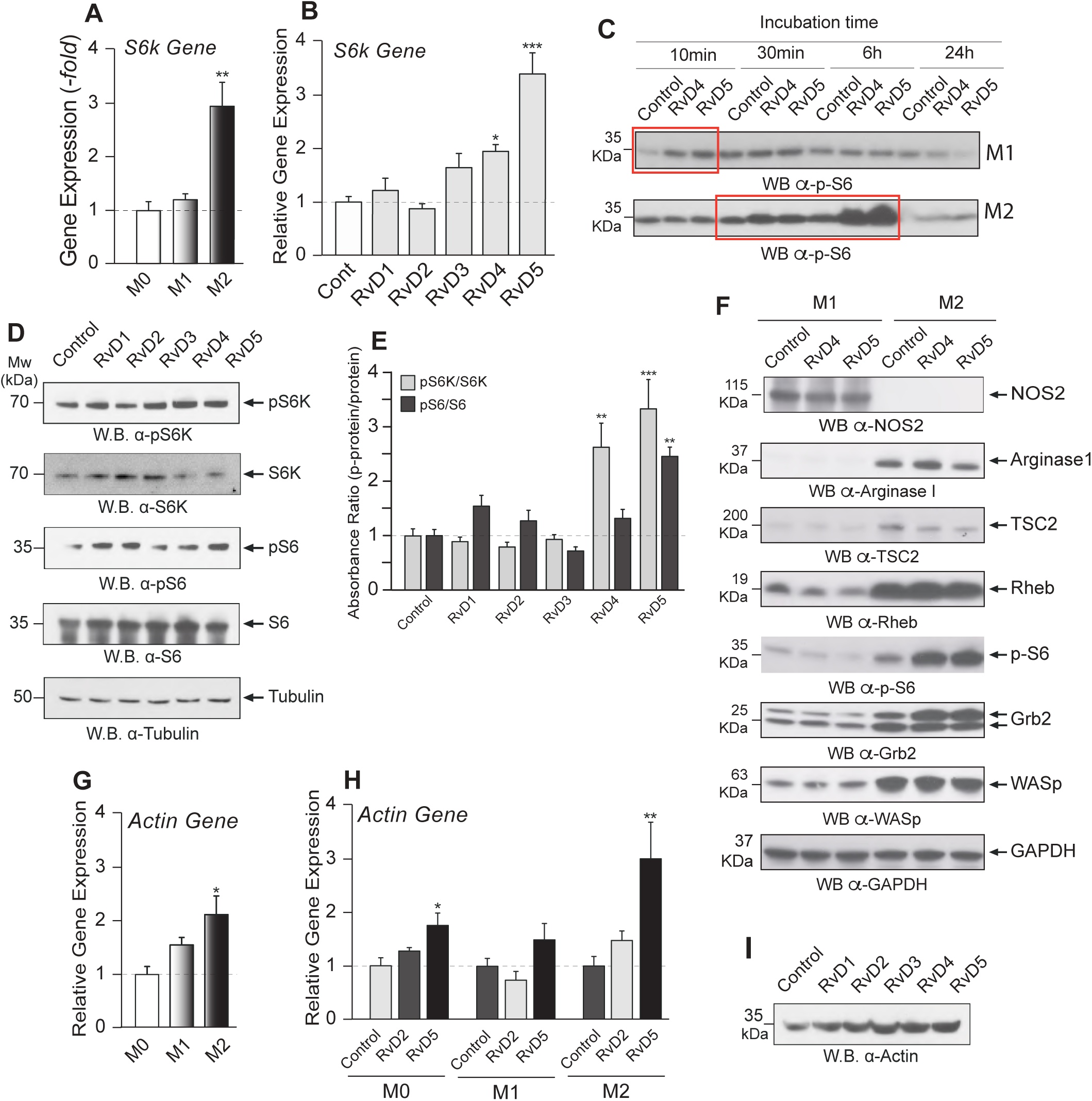
Mechanism for PLD-mediated actions of RvD5 in macrophages involving S6K and Actin. (A-E) S6K analysis. (A) Basal S6k gene expression by qRT-PCR in M0, M1, and M2 macrophages. (B) S6k gene expression changes upon 10 nM RvD1-5 treatment of M2 macrophages. (C) Effects of vehicle (Control), RvD4 or RvD5 (at 10 nM) on protein S6 phosphorylation [p(T232)-S6] in M1 or M2 macrophages. (D) Effects of RvD1-5 (at 10 nM) on S6K and phospho(T381)-S6K; S6 protein and phospho (T232)-S6 (with tubulin being loading control). (E) Densitometry of protein bands such as those shown in panel D, with the calculated ratio ‘phospho-protein’/’protein’ for S6K and for S6, in response to Resolvins-D1-5. (F) Analysis of proteins in the S6K signaling pathway (TSC2, Rheb and S6) and the Actin pathway necessary for phagocytosis (Grb2 and WASP). The top-most rows are NOS-2, which verifies the correct nature of M1 macrophages and Arginase-1, which verifies the correct nature of M2 macrophages. Bottom row: Loading controls with GAPDH. (G-I) Actin analysis. (G) Basal Actin gene expression (by qRT-PCR) in M0, M1, and M2 macrophages. (H) Actin gene expression in macrophages upon 10 nM RvD2 or RvD5 treatment for 6 hours in M0, M1, and M2 macrophages. (I) Actin protein levels in M2 macrophages upon 10 nM RvD1-5 treatment. Experiments from RAW264.7 macrophages are represented as mean ± SEM, n=3-4 independently. Statistical significance (*p < 0.05, **p < 0.01, and ***p < 0.001) was evaluated with one-way ANOVA and Tukey’s post hoc multiple comparisons.

We also observed a very large effect of RvD5 on S6 phosphorylation (**Fig. 7C**), a ribosomal protein that is a natural substrate of S6K. S6K and phospho (T381)-S6K; S6 protein and phospho (T232)-S6 are shown in Western blots (**Fig. 7D**). Densitometry of protein bands such as those shown in panel 8E, with the calculated ratio ‘phospho-protein’/’protein’ for S6K and for S6, in response to Resolvins-D1-5 (**Fig. 7E**) indicate the large effect of RvD4 and, especially RvD5.

Combining the pS6K pathway and WASp/Grb2 that our laboratory has shown (58) to be implicated in phagocytosis, **Fig. 7F** indicates that TSC2 is downregulated by RvD5 so it can no longer inhibit Rheb, which in fact is upregulated in M2 macrophages and, as a consequence, S6 phosphorylation is very robust. Even though RvD5 did not change the level of expression on WASP or Grb2, they both are expressed at high levels in M2 macrophages. PLD has been shown to be important for macrophage chemotaxis and phagocytosis of opsonized zymosan via PLD-Grb2-WASp heterotrimer formation and Actin polymerization (58). This could apply to efferocytosis, particularly since Actin gene expression is vastly increased in M2 (and so is S6K that can regulate phagocytosis) (**Figs. 7A and 7G**).

Moving next to studying a possible role of Actin polymerization as an effector of a PLD2-S6K pathway, basal Actin gene expression was measured in M0, M1, and M2 macrophages; with the basal level of Actin being the highest in M2 (**Fig. 7G**). Actin gene expression is further elevated with RvD5 treatment in M0 but, especially in M2 macrophages (**Fig. 7H**). Actin protein levels in M2 macrophages upon RvD1-5 treatment are shown by Western blot (**Fig. 7I**). Polymerization of Actin is needed for phagocytosis. All together these results suggest that at the protein level, PLD mediates actions of RvD5 on M2 macrophage functions by a PLD2-S6K-Actin signaling cascade.

### Proposed model summarizing the main findings

Schematic showing a proposed model of Resolvins D-series acting on M1-M2 conversion mediated by PLD2 signaling (**Fig. 8**). RvD5 reverses the inhibition of PLD2 by either PARN or by miR-138 or miR-887 thus increasing PLD2 expression and promoting PLD2 activity. This initiates PLD2 activation of S6K (we have found that RvD activates PLD in M2 macrophages that results in increased phospho-S6) and hence phagocytosis or efferocytosis and Actin that ultimately leads to CD206 cell surface expression contributing to M1 to M2 class switch (we have found that PLD induces macrophage polarization by affecting M2 markers and decreasing M1 markers). The M2 macrophages then carry out the key function of efferocytosis needed for resolution of inflammation.

**Figure 8.**
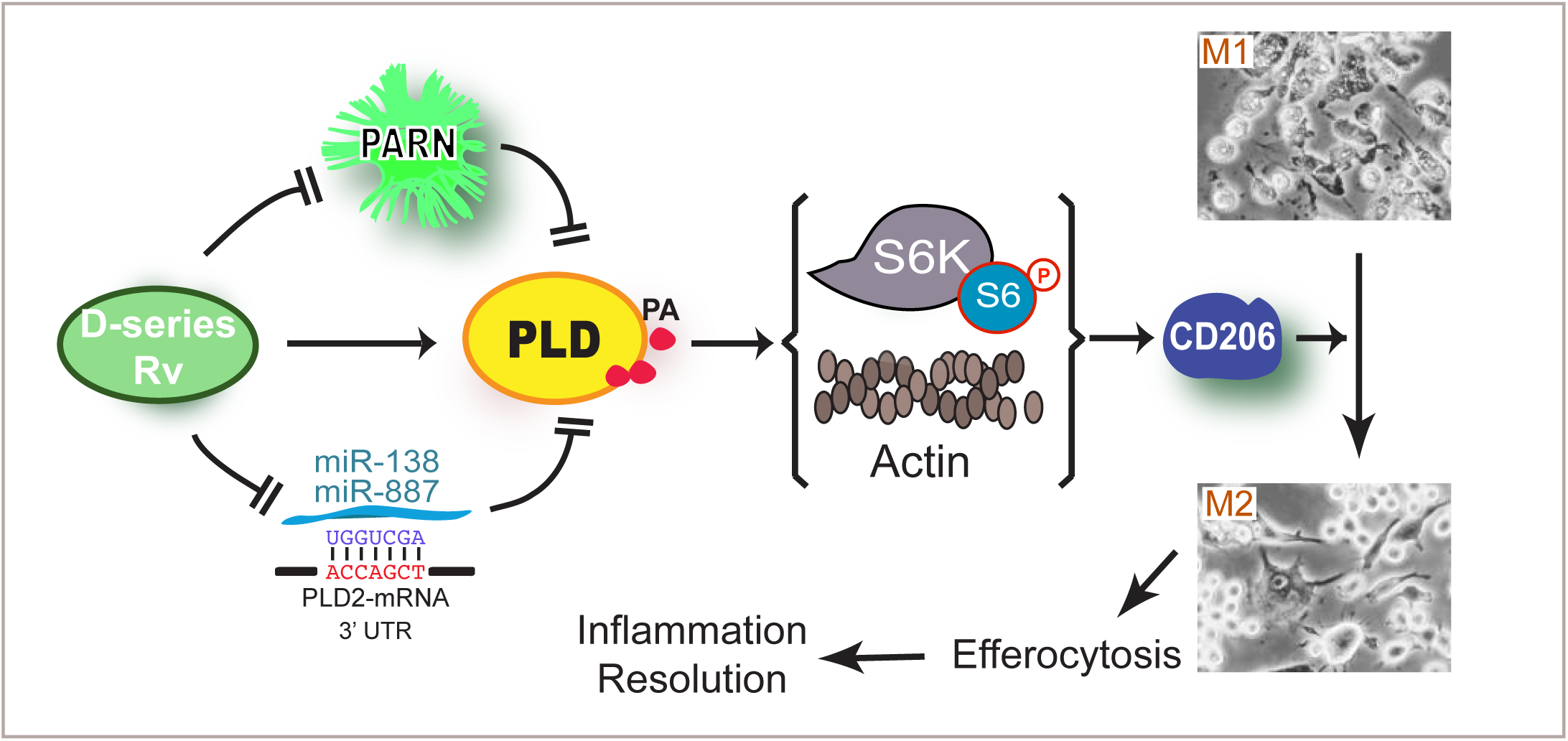
Proposed mechanisms of PLD-dependent RvD5-mediated role of macrophages in inflammation and resolution. A new model proposed to explain the intracellular signaling mechanisms of RvD5 action in macrophages as well as the RvD5-PLD2 cooperation in eliciting a class switch from pro-inflammatory M1 to pro-resolving M2 macrophages, wherein RvD5 reverses the inhibition of PLD2 by either PARN or by miR-138 or miR-887 thus increasing PLD2 expression and promoting PLD2 activity. This initiates PLD2 activation of S6K and Actin that ultimately leads to CD206 cell surface expression confirming to M1 to M2 class switch. The M2 macrophages then carry out the key function of efferocytosis needed for resolution of inflammation

## DISCUSSION

In view of the results presented in this study, PLD activity is high in M1 (inflammatory) and somewhat lower in M2 (scavengers) macrophages. Here, we show that Resolvins activate PLD, and it does so in both cell types. It appears that PLD is a necessary step to allow both chemotaxis and phagocytosis, regardless of the polarity of macrophages. An important control is the results in M’zero’ macrophages that are still “naïve”, and do not respond to Resolvins functionally in these settings. We indicate also in this study that macrophages switch polarity upon PLD2 transfection and several specific PLD-dependent pathways (miR, PARN) have been identified, which can explain the mechanism of resolution of inflammation for Resolvin, RvD5. To initiate the resolution phase of inflammation, apoptotic neutrophils have to be cleared (by M2 macrophages during efferocytosis), among other mechanisms. We have also found that Actin gene expression is high in M1 and even higher in M2, which could explain Actin dynamics thus allowing for continued cell/vesicle movement. This, along with the production of PA by PLD, could also be key for “membrane expansion” process that changes the morphology from M1 to M2 (the latter larger than the former).

Resolution of inflammation is a multi-step process and the resolvin-mediated inflammation resolution signaling mechanism is different in pro-inflammatory and anti-inflammatory macrophages exhibiting differential impact on PLD signaling. It is known that PLD is upregulated in activated neutrophils and macrophages during inflammation (59). Also, a role of lipoxins in regulating PLD mediated inflammation has been established, wherein lipoxins inhibit PLD activation by accumulation of pre-squalene diphosphate (PSDP) mainly in phagocytes (60). Even though FPR2 is regulated by PLD, whether or not the reverse is possible has not been tested. Since Resolvins act via GPCR (ALX/FPR2, GPR18, GPR32 or ChemR23) (53,61,62) we studied in this paper the role of Resolvins in regulating PLD protein, gene and functions. It has been shown that Resolvins increase during inflammation to promote resolution (51). As presented in Figure 1, Resolvin D5 in M1 and M2 macrophages is signaling via PLD2 suggesting a potential role in phosphorylation of S6 protein (Fig. 8), and protein synthesis with an increase in efferocytosis (Fig. 4).

RvD5 regulates inflammation resolution by promoting PLD2 protein expression, activity and PLD2-S6K signaling axis in M2 macrophages, leading to enhanced efferocytosis of apoptotic cells and thus inflammation resolution. This study presents a novel role of PLD in resolution of inflammation, which can be a basis for exploring the mechanism by which RvD5 upregulates PLD2 in M2 macrophages to promote efferocytosis.

Second-organ injury is a major concern that results from ischemia reperfusion. Lung is the second-organ most commonly affected from I/RI. Leukocytes mainly accumulate in the lungs during a second-organ injury exacerbating inflammation by releasing reactive oxygen species (ROS) and lysosomal enzymes. Myeloperoxidase (MPO) is the enzyme important for ROS generation and is a measure of neutrophil infiltration and activity. We report here that RvD5 could not reduce MPO activity in PLD2^-/-^ lungs with hind-limb ischemia reperfusion, suggesting PLD2 is important for RvD5’s pro-resolving activity. These results indicate that PLD2 intracellular signaling in M2 macrophages is a key signaling pathway in resolution of inflammation. This new information on the connection between PLD-RvDs may thus be used in human disease.

Macrophages are key phagocytes that along with dendritic cells, neutrophils and NK cells form the innate immune system. Macrophages can be polarized to either M1 or M2 depending on the environment and stimulus. M1 macrophages are pro-inflammatory and contain ROS- and proteolytic enzymes-rich granules. M1 macrophages secrete a number of pro-inflammatory mediators and their main function is kill and to phagocytose pathogens. On the other hand, M2 macrophages are anti-inflammatory and carry out efferocytosis of apoptotic cells (5,7,12). Herein, we demonstrate that PLD induces macrophage class-switch from M0 to M1-like and then M1 to M2-like by modulating cell surface marker expression and macrophage function. These findings suggest a time dependent role of PLD in the process of initiating inflammation and later in the resolution phase of inflammation.

## MATERIALS AND METHODS

### Materials

Human macrophages were derived from peripheral blood mononuclear cells (PBMCs) purchased from Boston Children’s Hospital Blood Bank. RAW264.7 murine macrophages (cat. # TIB-71) and DMEM (cat. # 30-2002) were obtained from ATCC (Manassas, VA, USA). RPMI 1640 with L-glutamine and 25 mM HEPES (cat. # SH30255.01) and ECL reagent (cat. # RPN2106) were from GE Healthcare Life Sciences (Logan, UT, USA). Fetal bovine serum (heat inactivated) (cat. # 900-108) and Penicillin/Streptomycin (10,000 units penicillin/10,000 mg/ml streptomycin) (cat. # 400-109) were from Gemini Bio-Products (West Sacramento, CA, USA). 5×10^−3^ M EDTA, pH 8.0 (cat. # 15575-038) was from Life Technologies (Carlsbad, CA, USA). Recombinant mouse M-CSF (cat. # 315-02) was from PeproTech (Rocky Hill, NK, USA). Sterile-filtered Histopaque 1119 (cat. #11191) was from Sigma-Aldrich (St. Louis, MO, USA). Mouse isotope control antibody (cat. # 553476) was obtained from BD Biosciences (San Diego, CA). shRNAs were purchased from Santa Cruz biotechnology (cat. # sh-44000-SH (shPLD1) and sh-44001-SH (shPLD2)). miR-138 (cat.# 002284) and miR-887 (cat.# 002374) were purchased from Taqman (cat.# 4427975). Resolvins were obtained from Dr. Charles Serhan’s lab and Avanti lipids (cat.# CAS# 810668-37-2 (RvD2) and CAS# 578008-43-2 (RvD5)).

### Animals

Animals used in the present study were 6-8 weeks old male or female wild-type C57BL/6 (Charles River Laboratories, Charleston, SC, USA), PLD1^-/-^ or PLD2^-/-^ mice (weighing 20-25 g). PLD1^-/-^ were obtained from Dr. Yasunori Kanaho’s laboratory, University of Tsukuba, Tennodai, Japan with exon 13 removed (63). PLD2^-/-^ were from Dr. Gilbert Di Paolo’s laboratory, Columbia University with exons 13-15 removed (64). The Pld1^-/-^ and Pld2^-/-^ mice were backcrossed with c57BL/6J mice for >7 generations (63,64). The mice were maintained in a temperature- and light-controlled environment with unrestricted access to food (laboratory standard rodent diet 5001 (Laboratory Rodent Diet 5001; PMI Nutrition International, St. Louis, MO, USA) containing 4.5% total fat with 0.3% ω-3 fatty acids and <0.02% C20:4 and were provided *ad libitum*) and water. The mice received veterinary care every day, and experiments were performed in accordance with the Harvard Medical School Standing Committee on Animals guidelines for animal care (Protocol 02570). Experiments for this manuscript have also followed the National Institutes of Health guide for the care and use of Laboratory animals (NIH Publications No. 8023, revised 1978).

### Ischemia-reperfusion-induced second-organ injury

Mice were anesthetized by i.p. injection of ketamine/xylazine mixture (80mg/kg;10mg/kg). Hind limb ischemia was induced using rubber band tourniquets tied on each hind limb (65). Hind limb ischemia was allowed for 1 h, after which the tourniquets were removed to begin reperfusion. Resolvin D5 (RvD5) was administered at 0.5 μg/mouse in vehicle (PBS + 0.1% ethanol) and compared to vehicle alone or no ischemia reperfusion control. RvD5 was administered intravenously ∼5 min prior to the start of reperfusion. At the end of this reperfusion period (2 h), these mice were euthanized with an overdose of anesthetic (isoflurane) and cervical dislocation. The lungs were quickly harvested and either flash frozen in liquid nitrogen and stored at −80°C or fixed and stored in 4% paraformaydehyde for H&E and immunohistochemistry. The right lungs (flash frozen) from individual mice were homogenized and centrifuged, and the tissue levels of myeloperoxidase (MPO) in the supernatants were determined using a mouse MPO ELISA (R&D systems, Biotechne, Minneapolis, MN, USA). The fixed tissues (left lungs) were sectioned and stained for hematoxylin and eosin (H&E) (AML laboratories Inc., FL, USA) to study its histology and degree of tissue damage.

### Myeloperoxidase (MPO) assay

MPO is an enzyme that is mostly present in neutrophils and correlates with the extent of neutrophil infiltration into tissues. The MPO assay was performed using the MPO Mouse Myeloperoxidase DuoSet ELISA (R&D systems, Minneapolis, MN) (cat# DY3667). Lungs were harvested from mice 2 hours after reperfusion. Samples were washed in cold phosphate-buffered saline (PBS, pH 7.4), immediately frozen on dry ice and stored at −80°C. For MPO assay, the lung samples were thawed, weighed, and homogenized in 1x PBS (pH 7.4) and centrifuged (1000 x g, 5 min, 4°C). The resulting supernatant was used for the MPO assay. Assay plates were prepared by coating the plate with capture antibody overnight at room temperature. The plates were then washed and blocked with blocking solution for 2h. After blocking, samples were added with respective control and standards. After thorough washing, detection antibody was added and incubated at room temperature for 2 h. Then the wells were washed thoroughly and streptavidin-HRP at 1:100 dilution was added for 20 minutes, followed by substrate for 20 min at room temperature. The reaction was stopped using a stop solution (2N H_2_SO_4_). The samples were read in a micro-plate reader at 450 nm with wavelength correction set to 540 nm or 570 nm. Results were expressed as units per microgram (wet weight) of protein in tissue.

### Histopathology of lung tissue

Lung tissue was fixed in 4% paraformaldehyde. The formalin fixed-tissue samples were used for sectioning for unstained sections or tissue histology H&E staining (AML Laboratory Inc., FL). EVOS™ XL Core Imaging System was used for microscopic analysis.

### Macrophage culture and polarization to M1 and M2 phenotypes

Human monocytes were isolated from de-identified peripheral blood *Leukopaks* obtained from Children’s Hospital Blood Bank (Boston, MA). Blood was obtained from healthy human volunteers giving informed consent under protocol #1999-P-001297 approved by the Partners Human Research Committee. All procedures were conducted in accordance with the relevant guidelines and regulations. The leukocytes-rich pack was used to isolate peripheral blood mononuclear cells (PBMCs) by Ficoll-Histopaque 1077-1 (Sigma-Aldrich, St. Louis, MO) density gradient. PBMCs were differentiated into macrophages (M0) using GM-CSF (20 ng/mL) or M-CSF (20 ng/mL) in RPMI culture media (Lonza, Walkersville, MD, USA) containing 10% FBS (Invitrogen, Carlsbad, CA, USA), 5 mM L-Glutamine (Lonza), and 1% penicillin and streptomycin (Lonza) for 7 days with media changes on days 3 and 6 at 37° C in an incubator with a humidified atmosphere of 5% CO_2_. For selected experiments (indicated as such in the figures’ legend) RAW264.7 murine macrophages were used instead of human macrophages. RAW264.7 cells were purchased from ATCC and maintained in DMEM (cat. # 30-2002) with 10% FBS and 1% penicillin and streptomycin at 37°C in an incubator with a humidified atmosphere of 5% CO_2_. For polarization of differentiated M0 macrophages towards M1 and M2 phenotypes, following published methods (45). Briefly, M’zero’ denoted M0 macrophages were polarized to M1 macrophages by the addition of 100 ng/mL LPS + 20 ng/mL IFN-γ to cultures and maintained for 1 day. M’zero’/M0 macrophages were polarized to M2 macrophages by the addition of 20 ng/mL IL-4 to cultures and maintained for 2 days. At the end of these time periods, M0, M1 or M2 macrophages were plated in a 6-well plate at 1×10^6^ cells/well or in 8-well chamber slides at 5×10^4^ cells/well overnight before experiments.

### Macrophage phagocytosis measurements

For phagocytosis, macrophages were incubated with fluorescent *E. coli*. and fluorescence was measured by real-time microscopy, unless otherwise indicated. Human macrophages were pre-incubated with vehicle (DPBS^Ca+/Mg+^) or D-series Rv (10 nM) in chamber slides kept in a Stage Top Incubation system for microscopes equipped with a built-in digital gas mixer and temperature regulator (TOKAI HIT model INUF-K14). Incubations were maintained for 15 minutes at 37 °C before addition of 1:50 green fluorescence protein (BacLightGFP)-labeled *E. coli* (5×10^8^ CFU/mL) (7 μg/mL, BacLight, Molecular Probes, Eugene, OR, USA). Fluorescence was assessed after 60 min (37 °C) using a BZ9000 real–time (BIOREVO) inverted fluorescence phase-contrast microscope (×20 objective) equipped with a monochrome/color-switching camera using BZ-II Viewer software (Keyence, Itasca, IL, USA). Incucyte™ Basic software computes green fluorescence intensity of phagocytosed *E. Coli* by macrophages from confocal images. In selected experiments (to directly compare phagocytosis with efferocytosis) phagocytosis of *E. Coli* was also measured by flow cytometry with 3 × 10^5^ cells treated with BacLight labeled *E. Coli* in the ratio of 1:50 cells:bacteria.

### Macrophage efferocytosis

For efferocytosis, macrophages were incubated with apoptotic human polymorphonuclear neutrophils (PMN). PMN were isolated by density-gradient Ficoll-Histopaque from human peripheral blood. Peripheral blood was obtained from healthy human volunteers giving informed consent under protocol # 1999-P-001297 approved by the Partners Human Research Committee. Isolated neutrophils were allowed to undergo apoptosis by plating 1 × 10^7^ cells/mL in 5 mL DPBS^Ca+/Mg+^ for 24 h in 100 mm × 20 mm Petri dishes and then stained with 10 μM, CellTrace™ CFDA, Molecular Probes, Eugene, OR, USA for 1h. Human macrophages were pre-incubated with vehicle (DPBS^Ca+/Mg+^) or D-series Rv (10 nM) for 15 minutes at 37 °C before addition of CFDA-labeled apoptotic PMNs (46,66). Apoptotic PMNs were added to 3×10^5^ macrophages in a ratio of 5:1. Apoptotic PMNs were added into 6-well plates with macrophages for 1 hour, after the cells were washed thoroughly and kept on ice, to stop any additional efferocytosis. These cells were harvested and subject to flow cytometry staining to measure mean fluorescence intensity as a read of efferocytosis with appropriate controls. A dilution 1:50 of Trypan blue was added to quench extracellular fluorescence.

### Cell culture and transfections

Human macrophages were transfected with 2 μg plasmid DNA, HA-PLD1 or myc-PLD2 for overexpression of PLD1 or PLD2 for 48 h or 300 ng shRNA, shPLD1 (cat. # sc-44000-SH) or shPLD2 (cat. # sc-44001-SH) (Santa Cruz Biotechnology, Dallas, TX) for silencing PLD1 or PLD2 for 72 h using JetPEI macrophage transfection reagent (cat. # 103-05N, Polyplus transfection, NY). RAW 264.7 macrophages were transfected with 1-2 μg plasmid DNA for 18-24 h, HA-PLD1 or myc-PLD2 using Amaxa® Cell Line Nucleofector® Kit V reagent (cat. # VCA-1003, Lonza, Switzerland).

### Flow cytometry

10^6^ cells in 100 μL FACS buffer were stained with fluorochrome-labelled antibodies for macrophage type-specific cell surface markers: anti-mouse CD23 (cat. # 101618, clone B3B4, Per.CP/Cy5.5, 7 μl, Biolegend, CA), anti-mouse CD38 (cat. # 102705, clone 90, FITC, 3 μl, Biolegend, San Diego, CA), anti-mouse CD80 (cat. # 104707, clone 16-10A1, PE, 3 μl, Biolegend, CA), anti-mouse CD206 (cat. # 141708, clone C068C2, APC, 3 μl, Biolegend, CA), anti-human CD80 (cat. # 305208, clone 2D10, PE, 10 μl, BD Pharmingen, CA), anti-human CD86 (cat. # 555660, clone 2331, APC, 10 μl, BD Pharmingen, CA), anti-human CD163 (cat. # 326512, clone RM3/1, PerCP-Cy5.5, 15 μl, BD Pharmingen, CA), anti-human CD206 (cat. # 551135, clone 19.2, FITC, 10 μl, BD Pharmingen, CA). Cell counts were obtained before flow staining by trypan blue exclusion and normalized to single stained and isotype controls on a FACSCanto II (BD Biosciences) or BD Accuri c6 (BD Biosciences) and FlowJo Ver. 10 software (Tree Star, Ashland, OR, USA) or BD FACS Xpress software were used for analyses.

### PLD activity

Total PLD activity was measured by an enzyme assay using [^3^H]-butanol in an *in vitro* transphosphatylation reaction to yield [^3^H]-phosphobutanol. Briefly, lysed macrophages (50 μg) or lysed lung tissue samples (50 μg) were processed for PLD2 activity in PC8 liposomes and [^3^H]n-butanol beginning with the addition of the following reagents (final concentrations): 3.5 mM PC8 phospholipid, 45 mM HEPES (pH 7.8), and 1.0 μCi [^3^H]n-butanol in a liposome form, as described (67). Samples were incubated for 20 min at 30°C with continuous shaking. Addition of 0.3 ml ice-cold chloroform/methanol (1:2) stopped the reactions. Lipids were isolated, dried (under N2) and suspended in chloroform:methanol (9:1) and then spotted on thin-layer chromatography plates along with 1,2-dipalmitoyl-sn-glycero-3-phosphobutanol (PBut) (Avanti polar lipids, Inc., AL) authentic standards. The amount of [^3^H]-phosphatidylbutanol ([^3^H]-PBut) that co-migrated with PBut standards (Rf∼0.45+0.36) was measured by scintillation spectrometry and background subtracted. Results were expressed as total PLD enzymatic activity as dpm/mg protein/min.

### Quantitative real-time Reverse Transcription Polymerase Chain Reaction (qRT-PCR) for mRNA measurements

Reverse transcription, coupled to qPCR, was performed following published technical details (68). Total RNA was isolated from macrophages with the RNeasy mini-kit following the manufacturer’s protocol (Qiagen, Valencia, CA, USA). RNA concentrations were determined using a Nano-Drop spectrophotometer, and samples were normalized to 29 ng RNA/μl. Reverse transcription was performed with 2 μg of RNA, 210 ng of random hexamers/primers, 500 μM dNTPs, 84 units of RNase OUT and 210 units of Moloney murine leukemia virus reverse transcriptase and incubated at 42°C for 55 min. Quantitative PCR reactions were run with 100 ng of total input RNA (3.45 μl), 10 μl of the gene expression assay (RT2 SYBR Green ROX qPCR Master Mix) (cat. # 330520, Qiagen, Valencia, CA, USA) and 1 μl of the relevant mouse RT2 qPCR Primer Assay. The following mouse primer sets were used from Qiagen (Valencia, CA, USA): TBP (PPM03560F) (used as housekeeping gene), PLD1 (PPH02835A), PLD2 (PPH02787A), S6K (PPH00791F), PLD3 (PPH02828A), PLD4 (PPH19319A) and PLD6 (PPH08933A), Actin (PPM02945B), PPARg (PPM05108C). Quantitative PCR conditions for the Stratagene Cycler were: 95°C for 10 min and then 50 cycles of the next 2 steps: 30 s 95°C and then 1 min 55°C, followed by 1 cycle of 1 min 95°C, 30 s 55°C and 30 s 95°C to establish the dissociation curves. The “cycle threshold” Ct values were chosen from the linear part of the PCR amplification curve where an increase in fluorescence can be detected at >10 S.E. above the background signal. ΔCt was calculated as: ΔCt = Avg. PLD Ct - Avg. Housekeeping Ct and gene-fold expression was calculated as: 2-(ΔΔCt) = 2^-(experimental condition ΔCt - control ΔCt).

### qRT-PCR for micro-RNA quantification

To measure miR-138 and miR-887 expression levels, cells that were polarized and treated with vehicle or Resolvins were used for RNA lysates using the Taqman micro-RNA Cells-to-CT kit according to the manufacturer’s protocol (Life Technologies; Cat. # 4391848). RNA concentrations were determined using a NanoDrop, and samples were normalized to ∼66 ng/μl RNA. Reverse transcription was performed in a 15 μl reaction volume with 1 μg of RNA, 1.5 μl 10T Buffer, 1 mM dNTPs, 3.8 units of RNase Inhibitor and 1 μl of Multiscribe Reverse Transcriptase and incubated in one-cycle at 16 °C for 30 min, 42 °C for 30 min and then 85 °C for 5 min. Quantitative PCR reactions were run in a 20 μl reaction volume using 10 μl Taqman Master Mix, ∼88 ng of total input RNA, 1 μl of the relevant micro-RNA gene expression assay (6-carboxyfluorescein (FAM)-labeled) multiplexed with U6 housekeeping gene. Taqman miR primers and fluorescent probes were from Life Technologies. Quantitative PCR conditions for the Stratagene Cycler were: 95 °C for 10 min and then 40 cycles of the next 3 steps: 15 s 95 °C and then 1 min 60 °C. Cycle threshold Ct values were obtained as indicated for mRNA qRT-PCR (above paragraph).

### SDS-PAGE and Western blot analyses

To confirm the presence of endogenous PLD1, PLD2 and S6K proteins in RAW264.7 macrophages, we performed SDS-PAGE and western blot analyses specific for each of these three proteins, as well as using TBP as the equal protein loading control. Approximately 150 μg of protein lysate (lysis buffer composition was: 50 mM HEPES, pH 7.2, 100 μM Na_3_VO_4_, 0.1% Triton X-100 and 1 mg/ml each of protease inhibitors (aprotonin and leupeptin) was loaded per each lane of the SDS-gels that were then used for Western blot analyses. For western-blot analyses, rabbit PLD1 (F-12) IgG (Santa Cruz Biotechnology, Santa Cruz, CA, USA) (cat. # sc-28314), rabbit PLD2 (N-term) IgG (Abgent, San Diego, CA, USA) (cat. # AP14669a), rabbit S6K IgG (49D7), S6 (cat. # 2217) and pS6 (cat. # 4858). (Cell Signaling Technology, Danvers, MA, USA) (cat. # 2708) and rabbit TBP IgG (Cell Signaling Technology) (cat. # 8515) were utilized as primary antibodies according to the manufacturers’ recommendations. Anti-rabbit or -mouse IgG HRP antibodies were from Cell Signaling Technology (cat. # 7074 and 7076, respectively). Immunoreactivities were detected using enhanced chemiluminescence (ECL) reagents from GE Heatlhcare (Pittsburgh, PA, USA) (cat. # RPN2106) and autoradiograph film.

### Statistical Analysis

The sample size for experiments was chosen empirically based on previous studies to ensure adequate statistical power. Data presented in the figures as bars are mean ± Standard Error of the Mean (SEM) (standard deviation/n^1/2^, were n is the sample size). Experiments were performed in technical triplicates (for qPCR measurements or for functional assays) or technical duplicates (for PLD activity assays or for flow cytometry) for n=3-5 independent experiments. Statistical significance (*p < 0.05, **p < 0.01, and ***p < 0.001) between means was assessed by two-tailed *t* test in case of experiments with only two groups (control and test). For multiple comparisons, one-way analysis of variance (ANOVA) with Tukey’s multiple comparison (or comparisons to controls) post hoc tests were used. Analyses of data were conducted using *GraphPad Prism 7* software (San Diego, CA).

## ACKNOWLEDGEMENTS

We thank Dr. Yasunori Kanaho and Dr. Gilbert Di Paolo for providing us the PLD2 and PLD1 knockout mice, respectively. We thank Dr. Kristen Fite for helpful suggestions and comments on this study. The following grants to Dr. Cambronero (J.G.C) have supported this work: HL056653-14 from the National Institutes of Health (NIH) and 13GRNT17230097 from the American Heart Association. Experiments in Dr. Serhan’s (CNS) lab were supported by National Institutes of Health grant P01GM095467.

This work is dedicated to the loving memory of our dearest colleague Dr. Julian Gomez-Cambronero.

## AUTHOR CONTRIBUTIONS

R.G. designed the study, carried out experiments and data analysis, prepared figures and wrote the manuscript. J.G.C. conceived and supervised research, contributed to manuscript preparation and critically edited the manuscript.

C.N.S supervised research and contributed to manuscript preparation and critically edited the manuscript. K.M.H. and K.S. carried out experiments and data analysis and contributed to manuscript and figure preparation. N.R.C contributed to data analysis and manuscript preparation. X.d.l.R. and S.L. performed and analyzed experiments and contributed to manuscript preparation.

## ABBREVIATIONS

Arg1: Arginase I
CD: Cluster of differentiation
DHA: Docosahexanoic acid
EPA: Eicosapentanoic acid
GPR32: G-protein coupled receptor 32
HLIR: Hind-limb ischemia reperfusion
IFNγ: Interferon gamma
IL-12: Interleukin 12
IL-4: Interleukin 4
iNOS2: Inducible Nitric oxide synthase 2
LPS: Lipopolysaccharide (Bacterial)
miR: micro RNA
MPO: Myeloperoxidase
PA: Phosphatidic acid
PARN: Poly(A)-tail ribonuclease
PBDM: Peripheral blood-derived macrophages
PC: Phosphatidyl choline
PLD: Phopsholipase D
PLD1-/-: Phospholipase D1 knockout
PLD2-/-: Phospholipase D2 knockout
PMN: Polymorphonuclear neutrophils
PPARγ: Peroxisome-proliferation antagonist receptor gamma
ROS: Reactive oxygen species
RvD1: 7S,8R,17S-trihydroxy-4Z,9E,11E,13Z,15E,19Z-docosahexaenoic acid/ Resolvin D1
RvD2: 7S,16R,17S-trihydroxy-4Z,8E,10Z,12E,14E,19Z-docosahexaenoic acid/ Resolvin D2
RvD3: 4S,11R,17S-trihydroxy-5Z,7E,9E,13Z,15E,19Z-docosahexaenoic acid/ Resolvin D3
RvD4: 4S,5,17S-trihydroxy-6E,8E,10E,13E,15Z,19Z-docosahexaenoic acid/ Resolvin D4
RvD5: 7S,17S-dihydroxy-4Z,8E,10Z,13Z,15E,19Z-docosahexaenoic acid/ Resolvin D5
S6: Ribosomal protein S6
phospho S6: Phosphorylated ribosomal protein S6
S6K: p70-S6 Kinase
SPMs: Specialized pro-resolving lipid mediators
TGF-β: Transforming growth factor beta
TNFα: Tumor necrosis factor alpha
WT: Wild-type

---

**Figure S1.**
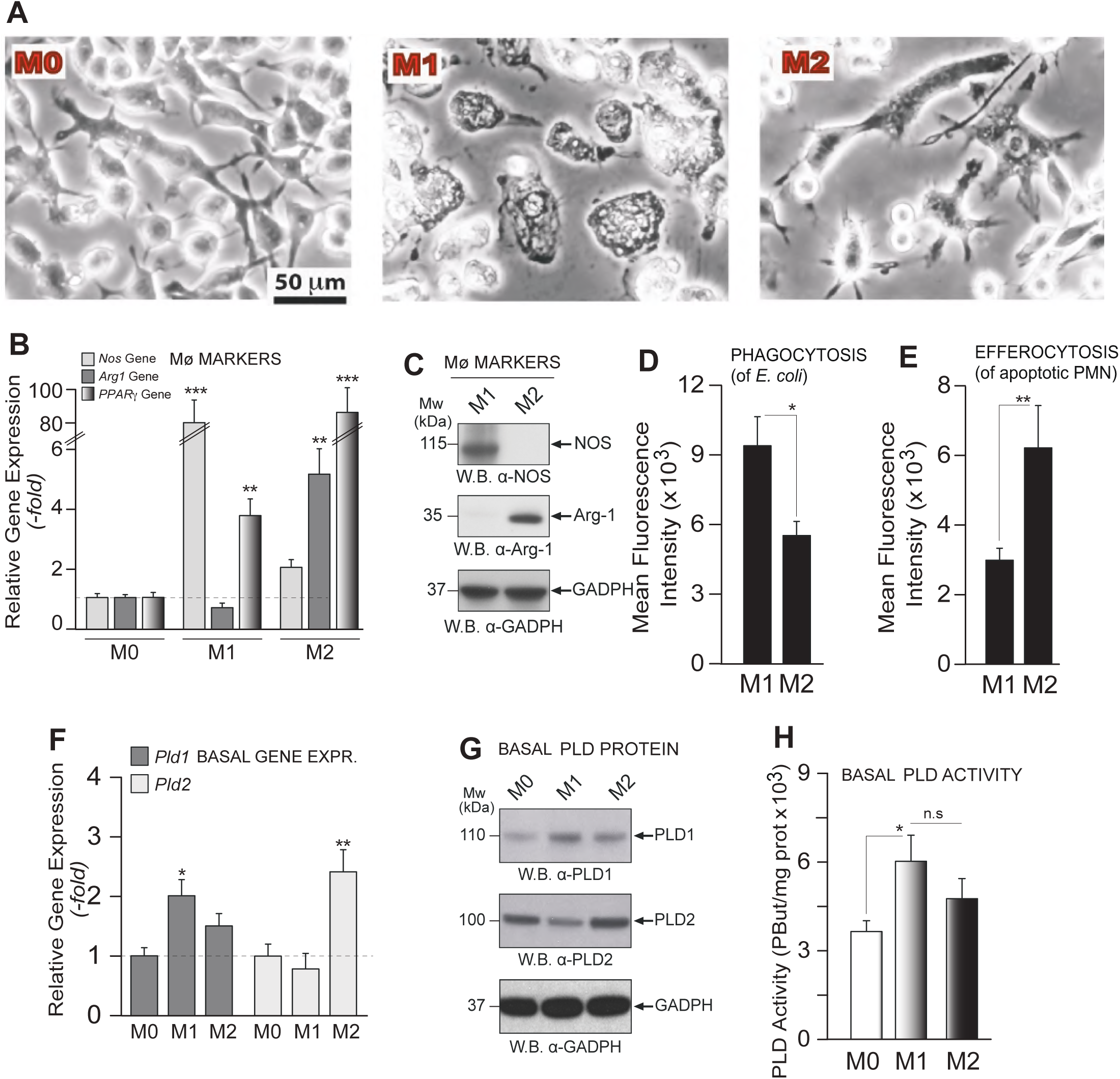
Assessment of gene, protein, activity, and functional levels in M1 and M2 macrophages. (A) Photomicrographs of representative morphology for M0, M1 and M2 macrophages by bright field microscopy. Results are representative of four experiments, and each panel is representative of 5 fields of view for each phenotype; scale bar = 50 μm. (B) Basal gene expression levels of iNos2, Arg1 and PPARγ by qRT-PCR. (C) Basal protein expression levels of iNOS2 and Arginase-1 in M1 and M2 macrophages by Western blot analysis (GAPDH are loading controls). (D) Phagocytosis. Basal levels of mean fluorescence intensity of phagocytosis of BacLightGFP-labeled E.coli by M1 or M2 macrophages measured by flow cytometry (to directly compare measurements with efferocytosis). E. coli was added to 3×105 macrophages in a ratio of 1:50 (macrophages:E.coli). (E) Efferocytosis. Basal levels of efferocytosis of green fluorescence intensity of green fluorescence (CellTraceTM-CFDA)-labeled apoptotic neutrophils (PMNs) by M1 or M2 macrophages measured by flow cytometry. Apoptotic PMNs were added in a ratio of 5:1 to 3×105 macrophages. (F) Basal gene expression levels of Pld1 and Pld2. (G) Protein expression of PLD1 and PLD2 by Western blotting (GAPDH are loading controls), and lipase activity (H) from M0, M1 and M2 macrophages. Data in bar graphs are mean ± SEM (n>3 independent experiments, each performed in technical duplicates, or triplicates for qRT-PCR and phagocytosis); statistical significance (*p < 0.05, **p < 0.01, and ***p < 0.001; n.s.=not significant) was evaluated with one-way ANOVA and Tukey’s post hoc comparing conditions to the M0 normalized controls for each panel (B,D,F), or two-tailed Student’s t test (G,H).

**Figure S2.**
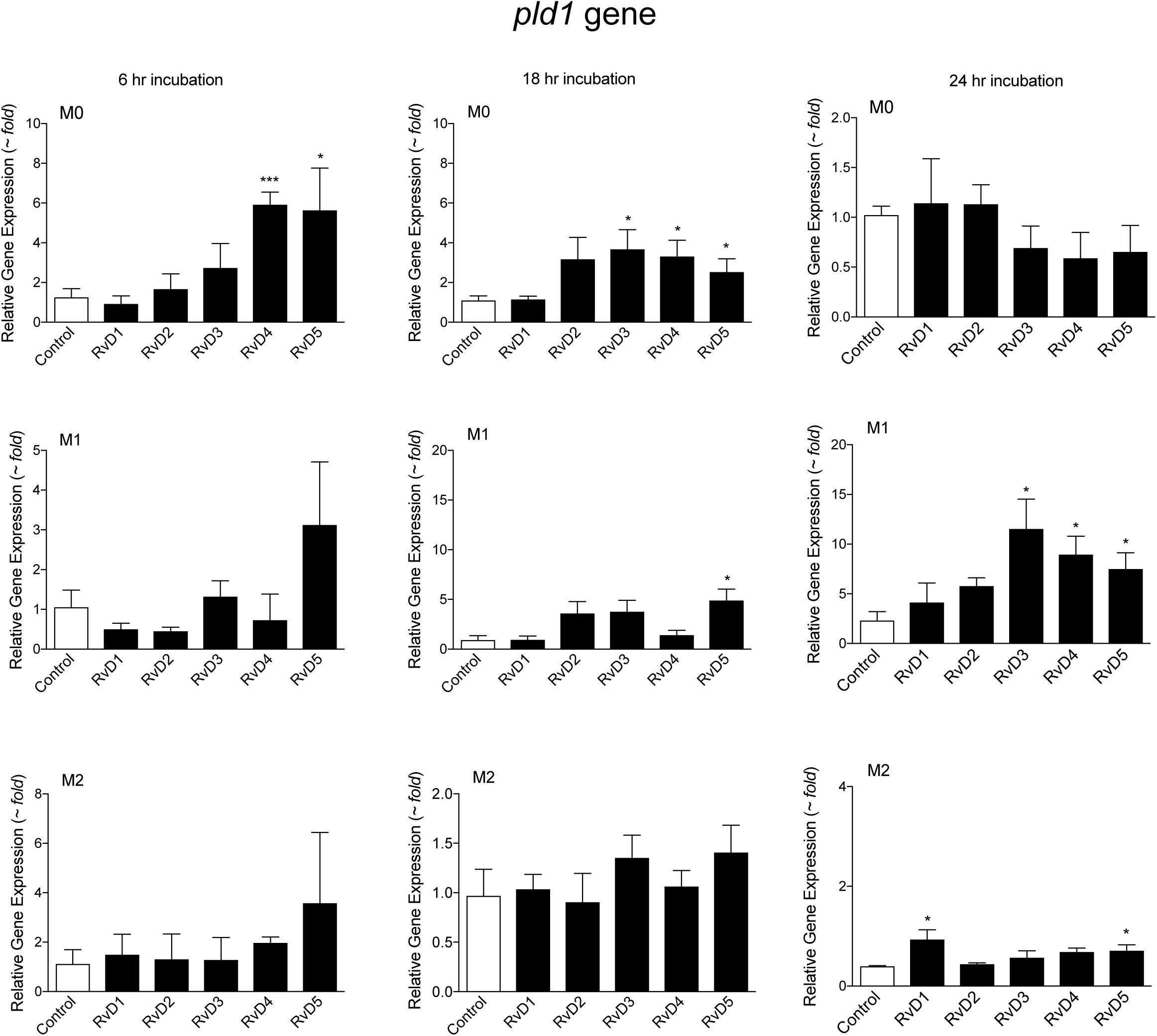
PLD1 gene expression in M1 and M2 macrophages time-course with RvD’s. See Main Figure 2 for specific Pld1 gene changes at 24 hours of M1 or M2 incubation with RvD’s.

**Figure S3.**
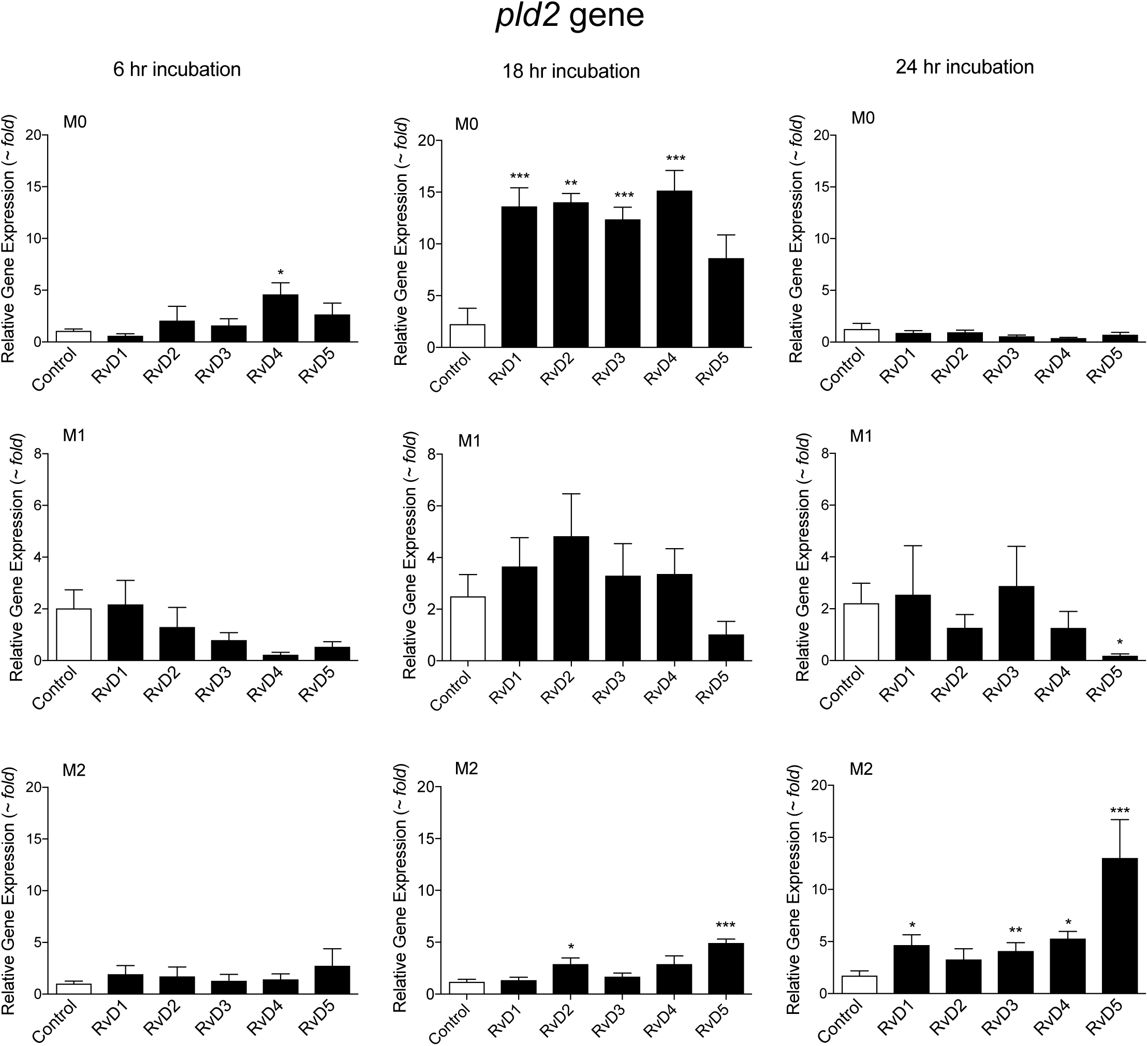
PLD2 gene expression in M0, M1 and M2 macrophages at different time lengths of macrophage incubation with RvDs. See Main Figure 2 for specific Pld2 gene changes at 24 hours of M1 or M2 incubation with RvDs.

**Figure S4.**
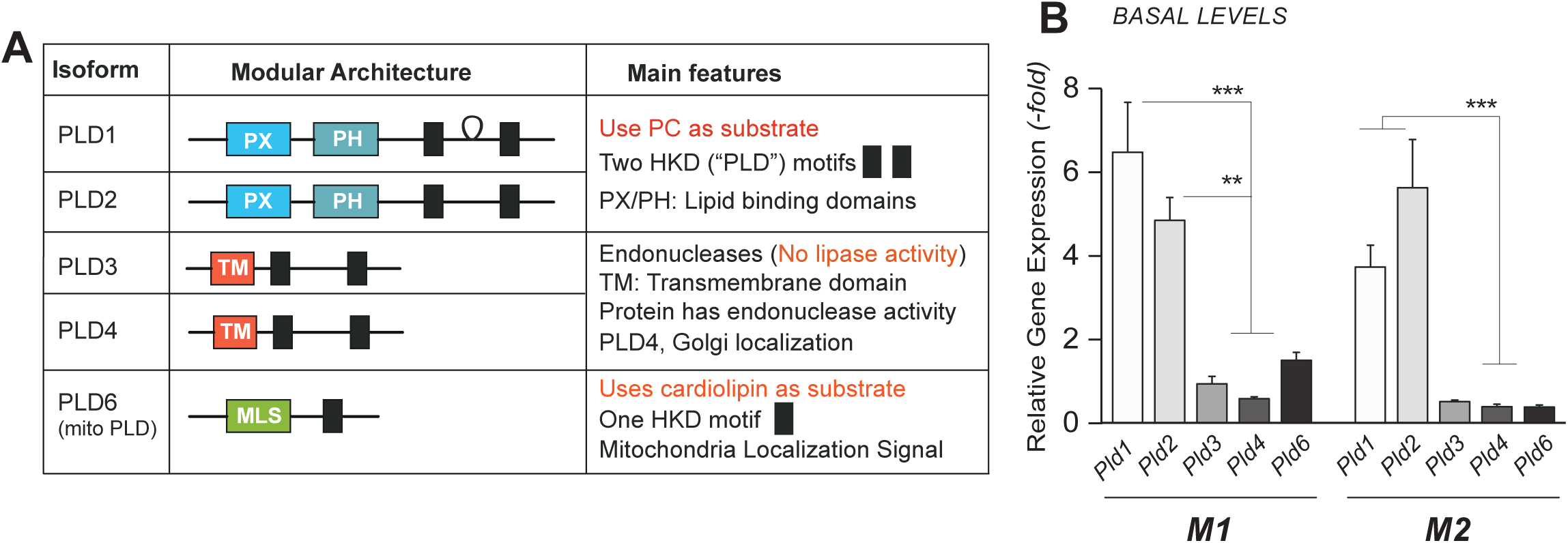
Classical and Non-classical PLD’s and their basal levels in M1 and M2 macrophages. (A) Scheme showing the molecular architecture and features of the “classical” (PLD1 and PLD2) and “non-classical” (PLD3, PLD4 and PLD6) mammalian PLD’s. HKD=catalytic site/lipase signature motif; PX=phox homology and PH=pleckstrin homology regulatory domains; or TM=transmembrane and MLS=mitochondria localization signal. (Note: PLD5 has not been fully described). (B) Comparison of basal gene expression levels between Pld1, Pld2, Pld3, Pld4, Pld6 using qRT-PCR in M1 and M2 macrophages.

**Figure S5.**
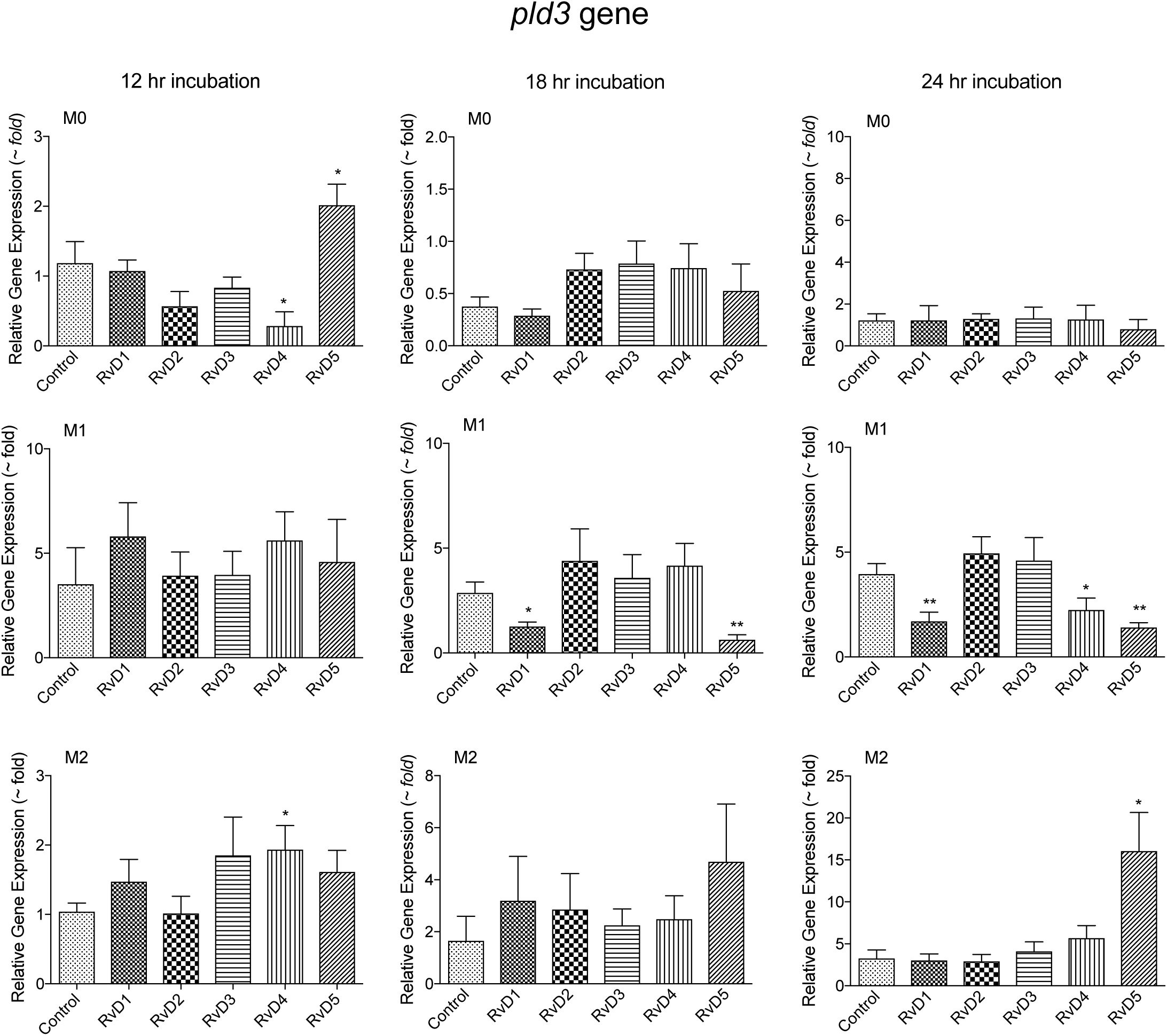
PLD3 gene expression in M0, M1 and M2 macrophages at different time lengths of macrophage incubation with RvDs. See Main Figure 3A for specific Pld3 gene changes at 24 hours of M1 or M2 incubation with RvDs.

**Figure S6.**
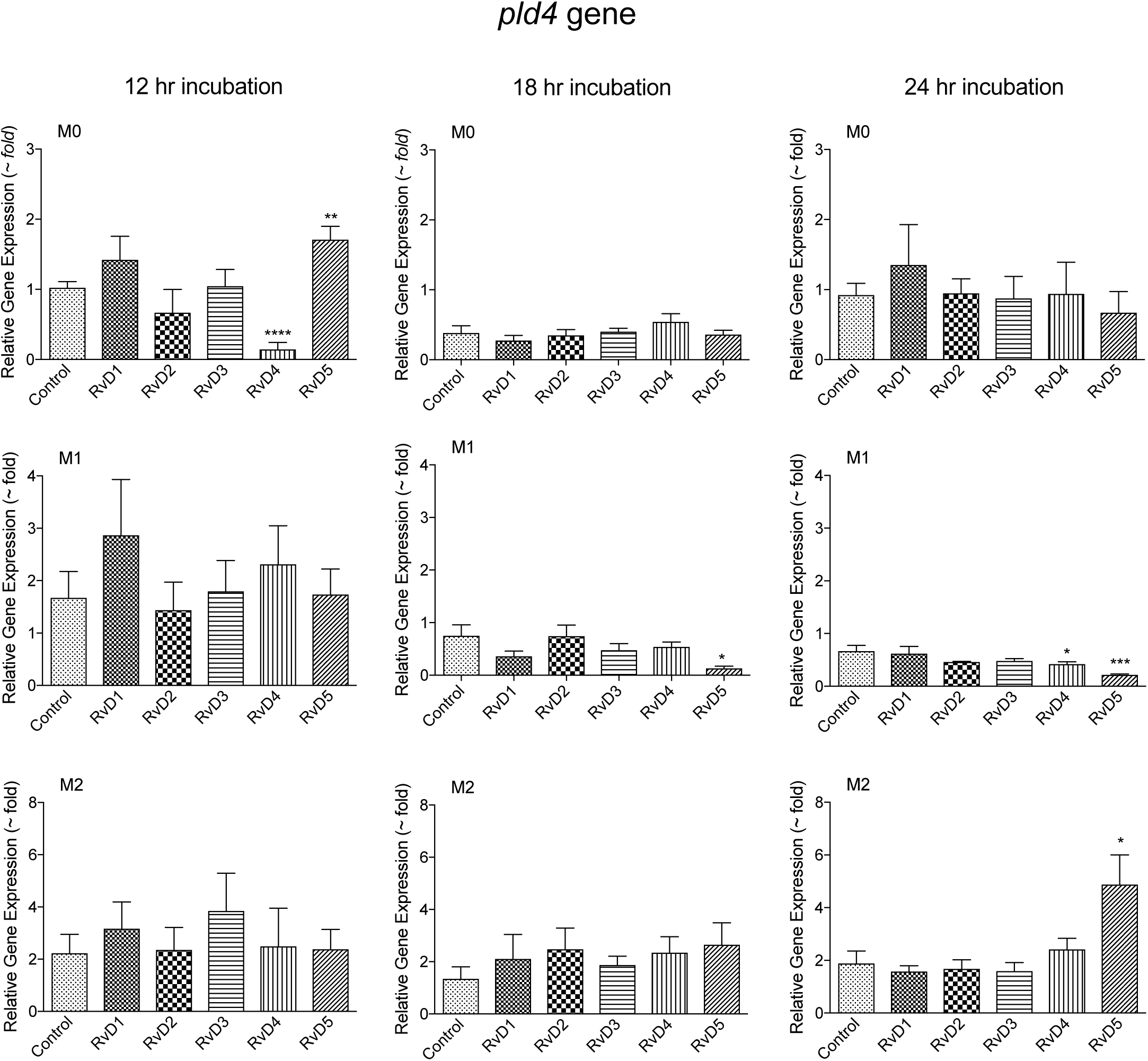
PLD4 gene expression in M0, M1 and M2 macrophages at different time lengths of macrophage incubation with RvDs. See Main Figure 3B for specific Pld4 gene changes at 24 hours of M1 or M2 incubation with RvDs.

**Figure S7.**
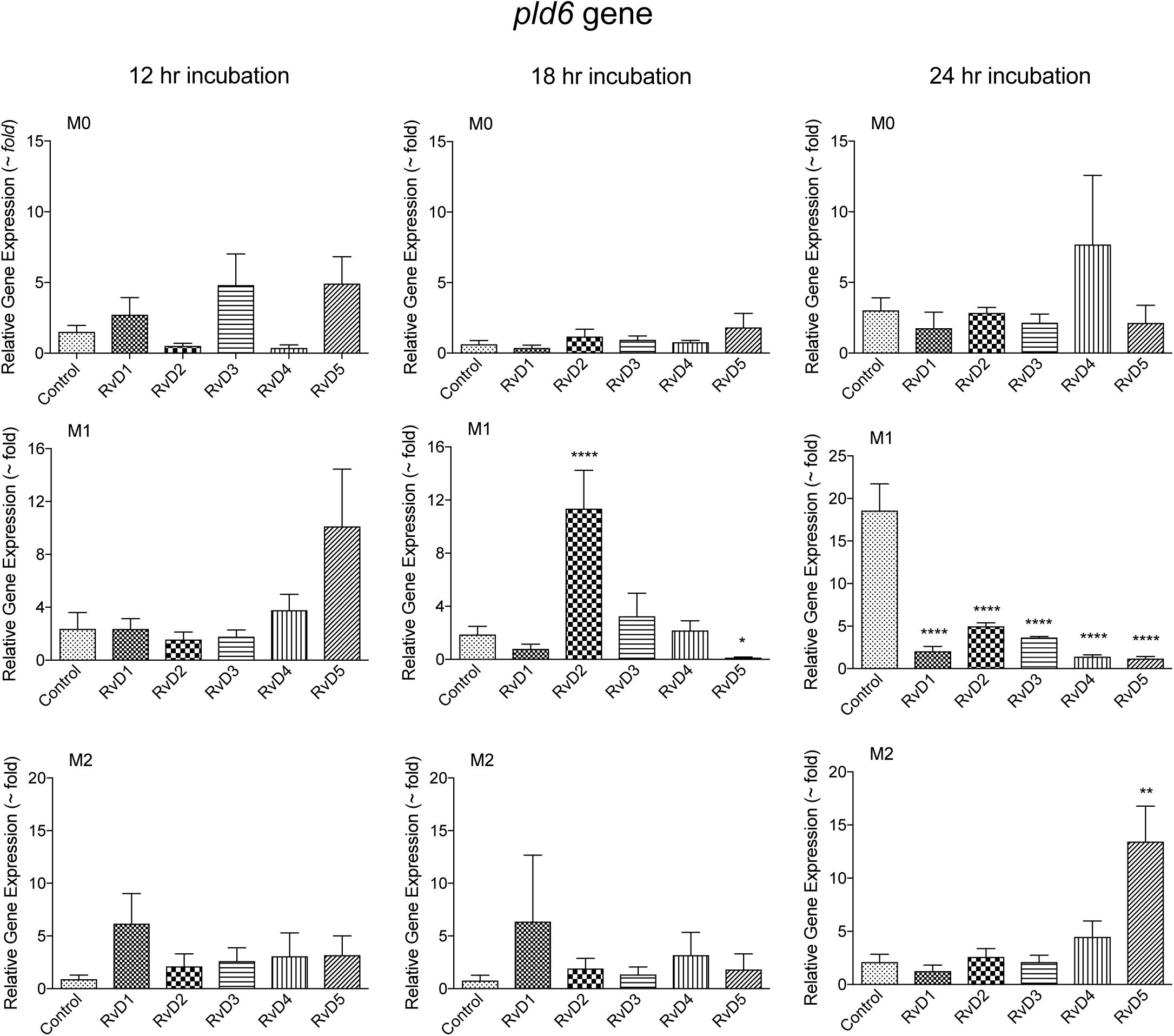
PLD6 gene expression in M0, M1 and M2 macrophages at different time lengths of macrophage incubation with RvDs. See Main Figure 3C for specific Pld6 gene changes at 24 hours of M1 or M2 incubation with RvDs.

**Figure S8.**
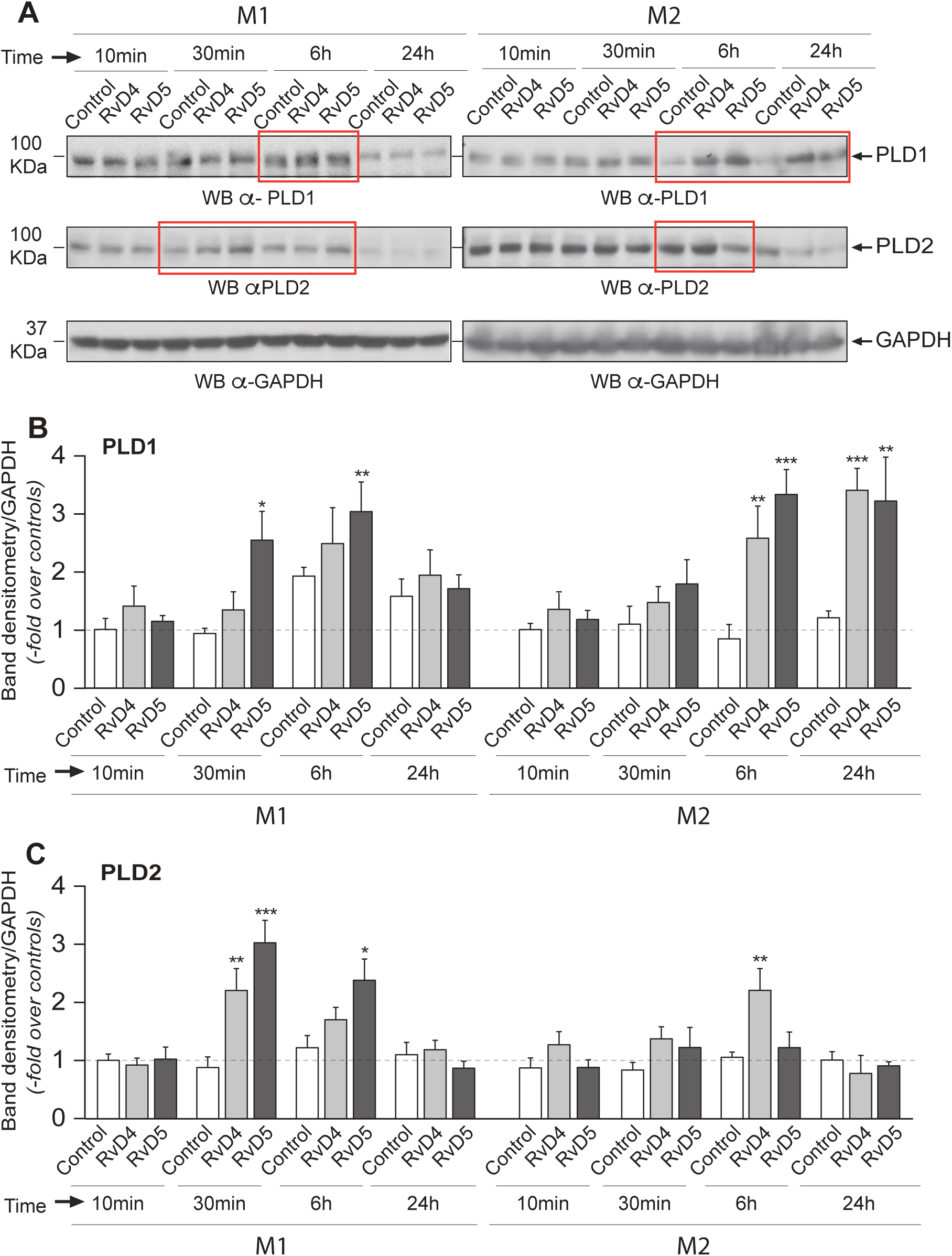
Effect of D-series Resolvins on PLD1 and PLD2 protein expression in M1 and M2 macrophages. (A) M1 or M2 macrophages were treated for 10 min, 30 min, 6 hours or 24 hours with vehicle (Control), 10 nM ResolvinD4 (RvD4) or 10 nM ResolvinD5 (RvD5). Cell lysates were obtained and used for SDS gel electrophoresis separation and Western Blotting, probed with either anti-PLD1 or anti-PLD2 antibodies (GAPDH as loading control). Red boxes highlight the times in which Resolvins had greater actions. (B) Quantification by densitometry scans of PLD1 protein bands in gels similar to the ones indicated in panel (A). (C) Quantification by densitometry scans of PLD2 protein bands in gels similar to the ones indicated in panel (A). Data is expressed as mean ± SEM from n=3 experiments; statistical significance (*p < 0.05, **p < 0.01, and ***p < 0.001) was evaluated with one-way ANOVA and Tukey’s post hoc comparing conditions to the controls (vehicle at 10 minutes) for each PLD and for each phenotype in panels B and C.

**Figure S9.**
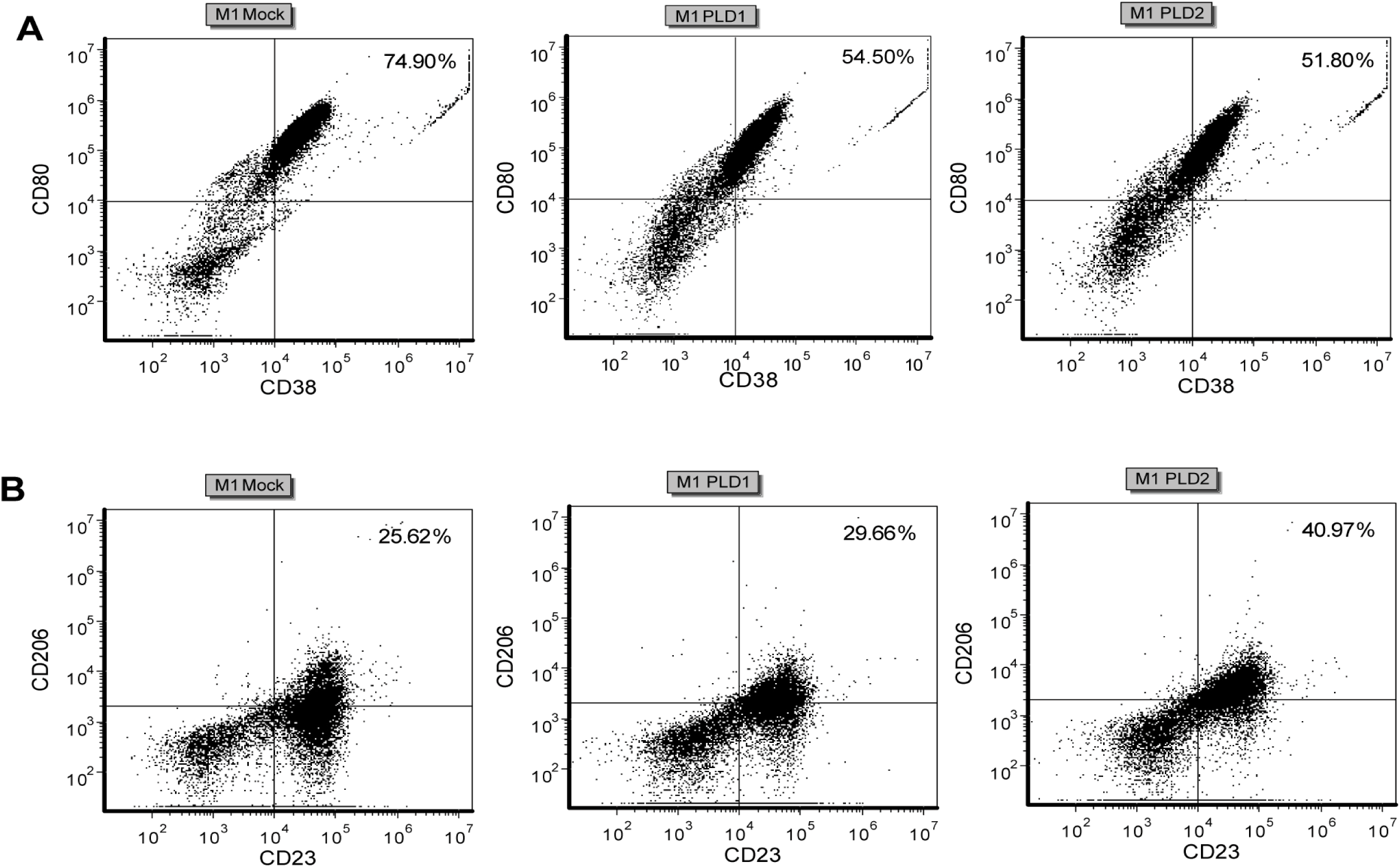
PLD ectopic expression causes macrophage polarization. See Main Figure 6D for plotting of this data.

